# A systemic clock brake: Period1 stabilizes the circadian network under environmental stress

**DOI:** 10.1101/2025.06.12.659230

**Authors:** Pureum Kim, Vinod Kumar, Nicholas Garner, Oliver Jayasingh, Gregg Roman, Shaun Walters, Tuan Vo, Quan Nguyen, Josephine Bowles, Trent Woodruff, Warrick Inder, Jeremy Hunt, Isabel Heyde, Henrik Oster, Oliver Rawashdeh

## Abstract

Precise alignment between internal circadian clocks and environmental light cycles is essential for physiological homeostasis and survival. However, the molecular mechanisms that preserve this synchrony across central and peripheral tissues remain poorly defined. Here, we uncover an unexpected role for the core clock gene *Period1 (Per1)* as a systemic modulator of circadian stability, regulating light-induced re-entrainment across the brain and body. In *Per1*-deficient mice, we show that loss of *Per1* accelerates clock realignment, influencing transcriptomic, metabolic, hormonal, and behavioral indicators of circadian realignment across multiple organ systems, including the suprachiasmatic nucleus (SCN) and peripheral tissues such as the liver, adipose tissue, and adrenal glands. Notably, this accelerated adaptation confers protection against jetlag-induced sleep disturbances, weight gain, and metabolic imbalance, underscoring a systemic role for *Per1* in maintaining circadian network stability. Mechanistically, unbiased spatial transcriptomics identified reduced expression of the arginine vasopressin (AVP), a key neuropeptide mediating SCN intercellular coupling, as the driver of circadian network instability. Weakened SCN synchrony permits enhanced flexibility of peripheral oscillator responses, expediting whole-body adaptation to shifted light-dark schedules. These findings position *Per1* as a critical regulator of circadian robustness, a buffer against light over-responsiveness, identifying a potential molecular target for mitigating circadian misalignment in contexts such as jetlag, shift work, and metabolic disease.

**Teaser:** What if beating jetlag was as simple as switching off a gene? Researchers show that disabling *Per1*, a core circadian regulator, accelerates body clock realignment and protects against sleep and metabolic disruption—highlighting new therapeutic possibilities for jetlag and shift work.

## Introduction

The rhythm of life itself is set by the 24-hour cycle of day and night—a relentless beat to which nearly all organisms have adapted by evolving internal clocks^1^ that synchronize with this cosmic metronome. In mammals, this system is orchestrated by the suprachiasmatic nucleus (SCN), a tiny cluster of neurons in the brain that aligns almost every cell in the body to the earth’s rotation^1^. At its core, this process is driven by an intricate transcription-translation feedback loop (TTFL), where proteins like Circadian Locomotor Output Cycles Kaput (CLOCK) and Brain and Muscle ARNT-Like 1 (BMAL1) spark a cascade of gene activity (e.g., *Period* [*Per1-3*] and *Cryptochrome* [*Cry1*, *Cry2*]) that culminates in the self-regulating oscillations^1–3^ we recognize as circadian (∼24 hour) rhythms^4,5^.

Light is the primary cue that keeps this internal clock ticking in sync with the environment—a delicate calibration that allows organisms to anticipate dawn and dusk with astonishing precision^6,7^.

However, when light exposure shifts abruptly, as in jetlag or shift work, the circadian system resists equally sudden adaptation^8,9^, creating a lag that disrupts physiology^8,10^. When chronic, this misalignment between internal and external time has been linked to serious health risks, including metabolic disorders, cardiovascular disease, cognitive decline, and type 2 diabetes^11–15^. While the SCN’s response to light is well-studied, the molecular mechanisms that regulate the rate of realignment—and thus the duration of jetlag—remain poorly understood.

The clock gene *Per1* is a known mediator of light-induced resetting in the SCN^7,16,17^, where it acts as an immediate-early gene in response to nighttime light exposure. However, its role in the broader circadian network, which includes peripheral clocks in organs such as the liver and adipose tissue, remains unclear. Traditionally, *Per1* has been thought to facilitate light-driven adaptation, but recent evidence suggests a more complex role.

Here, we hypothesized that *Per1* serves as a regulator of circadian network stability, acting to buffer the circadian system against excessive responsiveness to light. To test this, we used *Per1*-deficient mice and combined transcriptomic, metabolomic, and behavioral analyses to assess the impact of *Per1* loss across central and peripheral clocks. By mapping dynamic changes within the circadian network, we examined how *Per1* influences communication between the SCN and peripheral oscillators in response to environmental perturbation. Integrating spatial transcriptomics with physiological measures of circadian adaptation, we reveal a surprising twist: rather than facilitating circadian realignment, *Per1* functions as a brake, limiting the SCN’s responsiveness to sudden environmental changes.

This study challenges conventional views of *Per1* as a simple mediator of photoentrainment, instead positioning it as a critical modulator of circadian network stability with systemic impacts. These findings have broad implications for understanding circadian misalignment in jetlag, shift work, and metabolic disorders, providing a foundation for potential therapeutic interventions.

## Results

### *Period1* buffers light-induced circadian realignment of the sleep-wake rhythm

To determine whether *Per1* buffers the circadian system’s sensitivity to light and moderates phase shifts in behavioral rhythms, we first examined the phase-shifting capacity of a light pulse in the absence of *Per1*. Here we show that both *Per1*-proficient (WT) and *Per1*-deficient (*Per1^-/-^*) mice, when exposed to a 60-minute light pulse (35 lux) at circadian time (CT) 14, exhibited a delayed onset of wheel-running activity on subsequent days (Fig. S1*A*). However, in *Per1^-/-^* mice, the phase delaying effect of the CT14 light pulse was significantly amplified, resulting in a phase shift approximately three times larger than that measured in WT controls (WT: −1.16 ± 0.19 hours vs. *Per1^-/-^*: −3.79 ± 0.51 hours; *p=*0.0008, Student’s *t*-test; Fig. S1*A*, *C)*.

We next tested whether *Per1* similarly influenced the response to a phase-advancing light pulse at CT19. In contrast to CT14, there was no significant difference between genotypes (WT: 1.06 ± 0.29 hours vs. *Per1^-/-^*: 0.96 ± 0.49 hours; *p=*0.65, Student’s *t*-test) (Fig. S1*B*, *C*), suggesting that *Per1* selectively modulates phase-delay responses. This directional asymmetry in light sensitivity led us to hypothesize that *Per1* selectively buffers the circadian oscillator’s response to light—challenging its widely accepted role as a general facilitator of light-driven circadian entrainment.

The enhanced phase-delaying response in *Per1*^−/−^ mice may be attributed to three mechanisms: (a) altered phase-shifting capacity via increased wheel-running activity, which could enhance serotonergic input to the SCN and phase shift the clock ^18^; (b) exercise-facilitated entrainment through sleep homeostasis, strengthening sleep pressure’s influence on the circadian clock ^19,20^; or (c) a novel role for *Per1* in buffering, rather than facilitating, the SCN’s response to light. However, our findings do not support the first two explanations, as activity levels were similar across genotypes (Fig. S2).

This led us to focus on the third possibility: that *Per1* buffers the circadian oscillator’s response to light—challenging its widely accepted role in promoting light-driven circadian entrainment. If this is the case, the enhanced phase-delaying response at CT14 predicts that the entrainment of *Per1*-deficient mice to a phase-delayed light-dark (LD) cycle is accelerated, while the absence of a response to a CT19 light pulse indicates that phase advancing the LD cycle, predicts that *Per1* deficiency would have minimal impact on the dynamics of phase-advancing entrainment. To test this hypothesis, we subjected *Per1^-/-^*and WT mice to a reversed LD cycle as shown in Fig. 1*A* – a jetlag protocol simulating travel across 12 time zones – and measured the rate by which the sleep-wake cycle realigned. The sleep-wake cycle of WT mice exhibited an average daily shift of 1.75 ± 0.21 hours, with a PS_50_ (the time required for a 50% re-alignment to the shifted LD schedule) of 3.01 ± 0.39 days (Fig. 1*B*). Remarkably, *Per1^-/-^*mice realigned their sleep-wake rhythms nearly five times faster, with an average daily shift of 5.87 ± 0.16 hours and a PS_50_ of 0.73 ± 0.11 days, near-immediate entrainment (PS_50_ WT vs *Per1^-/-^*, *p<*0.0001, student’s *t*-test; Fig. 1*B*).

**Fig. 1.**
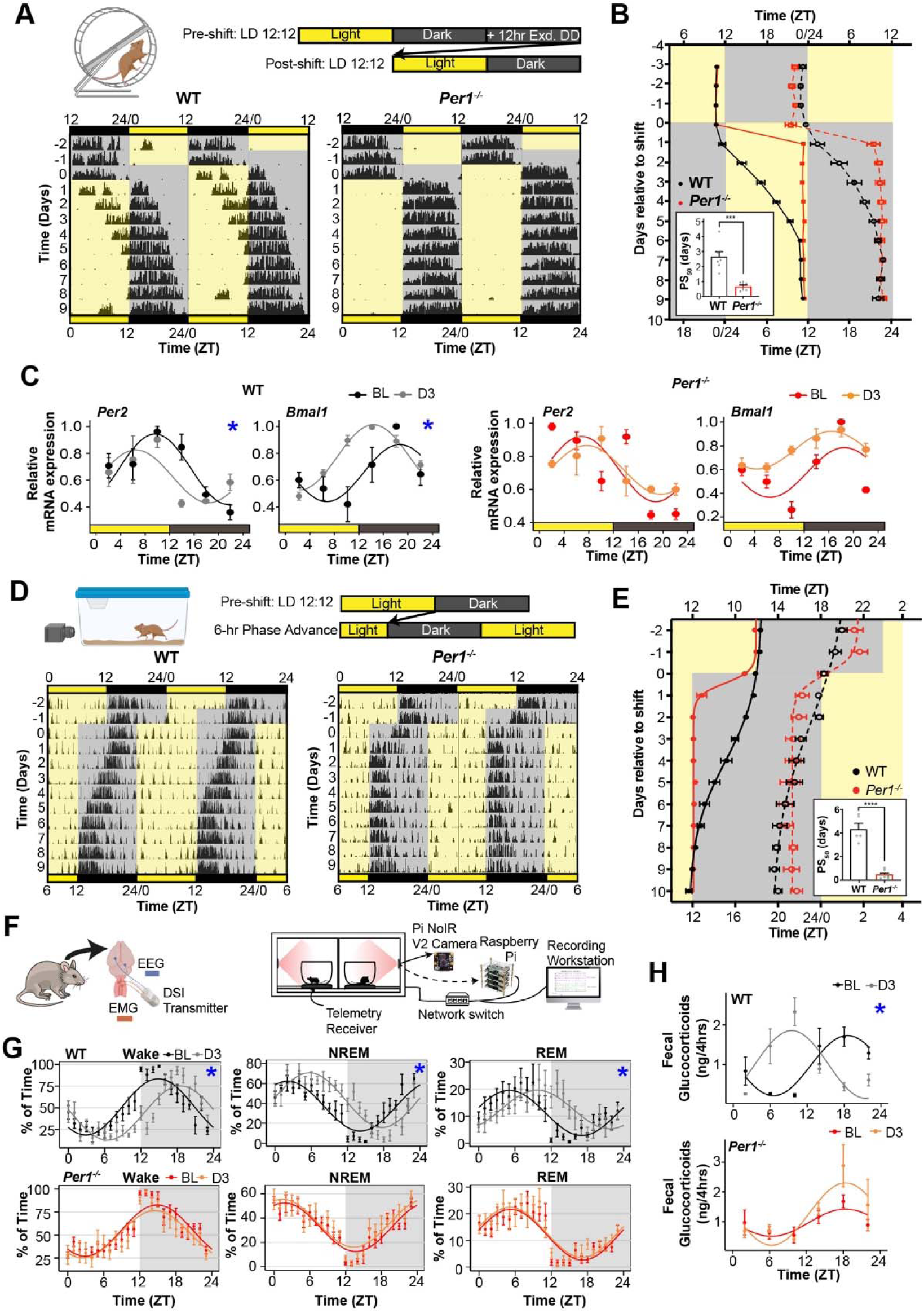
*Period1* restricts light-induced phase shifts. (**A**) (Top) Experimental timeline: Mice were initially entrained to a 12:12 light-dark (LD) cycle, then subjected to a 12-hour phase delay. (Bottom) Representative double-plotted actograms show running wheel activity before and after the phase delay, with pre- and post-shift zeitgeber times (ZTs) indicated. (**B**) Onset (solid line) and offset (dashed line) of running activity before and after the 12-hour phase delay. Phase shift (PS_50_) values for entrainment in wild-type (WT) and *Per1^-/-^* mice, with significant differences noted (*p*<0.0001, Student’s *t*-test; WT, n=6; *Per1^-/-^*, n=5). (**C**) SCN expression profiles of *Per2* and *Bmal1* on day 3 post shift (D3) versus baseline (BL) in WT and *Per1^-/-^* mice (WT BL vs. D3: *Per2*, *p*=0.021; *Bmal1*, *p*=0.0041, CircaCompare). (**D**) Representative actograms of locomotor activity in WT and *Per1^-/-^* mice before and after a 6-hour phase advance. (**E**) Onset (solid line) and offset (dashed line) of activity following a 6-hour phase advance, with PS_50_ comparison between genotypes (*p*<0.0001, Student’s *t*-test; WT, n=6; *Per1^-/-^*, n=5). (**F**) Schematic of telemetry transmitter placement for EEG recording in brain and trapezius muscle to assess sleep. (**G**) Rhythms of Wake, NREM, and REM sleep fitted with cosinor curves for WT and *Per1^-/-^*mice on D3 post 6-hour phase advance compared to BL (WT BL vs. D3: Wake, *p*<0.05; NREM, *p*<0.05; REM, *p*<0.05 and *Per1^-/-^* BL vs. D3: Wake, *p>*0.05; NREM, *p*>0.05; REM, *p*>0.05; CircaCompare). (**H**) Glucocorticoid rhythms from fecal samples on D3 post 12-hour phase delay, compared to BL. Data presented are mean ± SEM; statistical comparisons to baseline unless noted otherwise. \**p*<0.05, \*\*\**p*<0.0005.

Although light is the primary zeitgeber for circadian entrainment, acute light exposure can also modulate arousal states independently of the circadian system. In nocturnal rodents, light exposure can induce sleep, a phenomenon known as negative masking or photosomnolence, which can complicate running wheel activity-based assessments of circadian phase; wheel-running actigraphy do not capture off-wheel activity. To verify the accelerated entrainment in *Per1^−/−^* mice, we repeated the protocol using video-based monitoring of sleep-wake rhythms across three cycles of reversed LD exposure. Consistent with the running-wheel activity data in Fig.S1*D*, only *Per1^−/−^*mice achieved full entrainment within the recording timeframe (Fig. S1*D*).

Accelerated entrainment was also observed at the molecular level within the SCN. Within three days of LD reversal, the molecular clock components *Per2* (negative element) and *Bmal1* (positive element) were fully entrained in the SCN of *Per1^−/−^* mice (Figs. 1*C*, S3). In contrast, WT SCNs had not achieved full entrainment by day 3 (D3). Increased temporal resolution, including SCN sampling on day 1 (D1) post-shift, revealed that *Per2* rhythms in *Per1^−/−^* SCNs phase-aligned almost immediately with the 12-hour shift (Δphase BL vs. D1*, Per2 p*=0.35; *GLMMcosinor*), and *Bmal1* was fully entrained by D3 (Δphase BL vs. D3*, Bmal1 p*=0.13; *GLMMcosinor*) (Fig. S3). These data indicate that daily phase adjustments in SCN clock gene expression rhythms are significantly larger in *Per1^−/−^*mice than initially predicted based on D3 and D5 measurements alone.

Collectively, our findings suggest that the *Per1* protein (PER1) functions as a buffer, moderating light-dependent entrainment of the SCN molecular oscillator and downstream clock-regulated sleep-wake rhythms.

Having established that the absence of *Per1* amplifies the circadian clock’s response to phase-delaying light cues, we next investigated whether this accelerated realignment extends to phase advances, thereby addressing its role in bidirectional entrainment.

### *Period1* deletion accelerates bidirectional entrainment

While a substantial body of literature highlights a pivotal role for *Per1* in light-induced phase shifting ^21–24^, our findings challenge the notion that *Per1* is required for photic entrainment. Instead, we show that *Per1* is dispensable for light-mediated circadian adjustment and acts to inhibit rapid adaptation to shifts in the light-dark (LD) environment.

Despite similar responses of *Per1^-/-^* and WT to a phase-advancing light pulse at CT19 (Fig. S1*A - C*), the absence of *Per1* leads to accelerated entrainment to both phase-delaying and phase-advancing shifts in the 12:12 LD cycle (Fig. 1*D*, *E*). WT mice exhibit significantly slower entrainment than *Per1^-/-^* mice to a full 12-hour LD reversal and to a phase advance (PS_50_ WT vs *Per1^-/-^*, *p<*0.0001, student’s *t*-test; Fig. 1*D*, *E*). These results suggest that the acute behavioral response to a light pulse does not necessarily predict the rate of behavioral entrainment to an LD shift.

The robust bidirectional accelerated behavioral realignment in *Per1*-deficient mice raises the question of whether this faster realignment also extends to physiological processes tightly regulated by circadian clocks, such as sleep and hormone rhythms. To explore this, we examined the re-entrainment dynamics of sleep states and key systemic hormones.

### *Period1* gates the entrainment of sleep and peripheral sleep-modulating hormones

Recognizing that activity data from running wheels and home cages do not directly measure sleep-wake states, we assessed entrainment for rhythms in vigilance states (sleep and wake) to a 6-hour phase advance in the LD cycle using electroencephalography (EEG) (Fig. 1*F*, *G*). By day 3 post-shift, wakefulness, non-rapid eye movement (NREM) sleep, and rapid eye movement (REM) sleep rhythms in *Per1-/-* mice had qualitatively re-aligned with the pre-shift baseline (D0) (Fig. 1*G* and Table 1).

**Table 1.** Summary table of *CircaCompare* statistical comparison for vigilant states of sleep and glucocorticoids between the BL and D3 post phase shift for each genotype. Statistical significance is determined from comparison between baseline and post shift days within each genotype. * indicates *p*<0.05.

| Peak Phase (ZT) | WT |  | *Per1*<sup>-/-</sup> |  |
| --- | --- | --- | --- | --- |
|  | BL | D3 | BL | D3 |
| Wake | 14.88 | *18.51 | 14.88 | 14.54 |
| NREM | 2.09 | *5.69 | 1.73 | 1.39 |
| REM | 5.07 | *9.69 | 5.21 | 5.09 |
| Glucocorticoids | 18.55 | *9.49 | 18.34 | 18.36 |

Sleep and wake reflect distinct physiological states governed by both circadian and homeostatic mechanisms, heavily modulated by rhythmic hormones such as melatonin and glucocorticoids (GCs) ^25^ ^26,27^. Thus, the rapid realignment of the sleep-wake rhythm in *Per1^-/-^*mice indicates accelerated entrainment of systemic physiological processes. In line with this, GC rhythms in *Per1^-/-^* mice fully re-entrained to the new LD cycle within three days (Δphase BL vs. D3 = 0 hours, *p*>0.05) (Fig. 1*H* and Table 1), which aligns with the accelerated entrainment of the sleep and wake cycles. In contrast, WT mice showed a significantly delayed GC re-entrainment (Δphase BL vs. D3 = 9.06 hours, *p*<0.0001).

### *Period1* as a gatekeeper of peripheral circadian clocks

Our observations of rapid sleep and hormonal realignment led us to investigate whether *Per1*’s buffering effect extends to the resetting of circadian oscillators in peripheral organs. To further elucidate *Per1*’s role in circadian entrainment, we measured the phase-shifting dynamics in the rhythmic expression of the clock genes *Per2* and *Bmal1* in the liver and white adipose tissue (WAT), two metabolically active organs. Under baseline (entrained) conditions, the phase and amplitude of *Per2* and *Bmal1* rhythms were comparable between WT and *Per1*^-/-^ mice (Fig. *2A–E*), with both tissues displaying delayed phase profiles relative to the suprachiasmatic nucleus (SCN). Consistent with previous findings in WT mice ^26,28,29^, peripheral clock gene entrainment lagged behind the SCN (Fig. 2*B*-*E*).

**Fig. 2.**
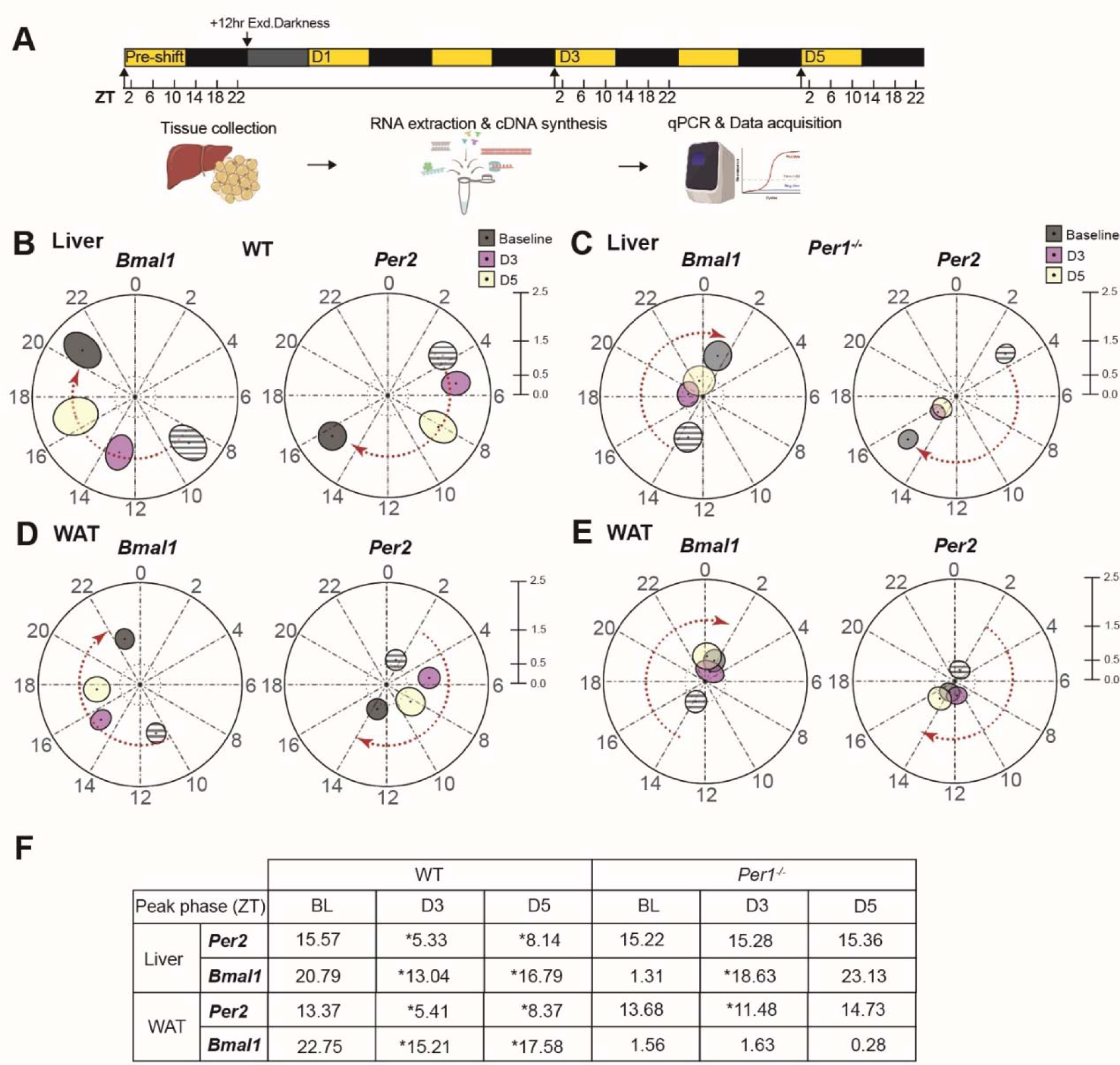
*Period1* modulates re-entrainment of peripheral clocks. (**A**) Experimental protocol: 12-hour phase shift with tissue collections in 4-hour intervals pre-shift and on day 3 (D3) and day 5 (D5) post-shift. (**B**–**E**) Polar plots of *Bmal1* and *Per2* expression rhythms in liver (**B, C**) and white adipose tissue (WAT) (**D, E**) of WT and *Per1^-/-^* mice at baseline and on D3 and D5 post-shift. Peak phase in zeitgeber time (ZT) is plotted as a circular shape, with amplitude represented by the distance from the center. Arrows indicate re-entrainment direction. Hashed grey circles denote baseline rhythms from pre-shift conditions. (**F**) Summary table of *Bmal1* and *Per2* peak phases in liver and WAT of WT and *Per1^-/-^*mice on D3 and D5, with baseline values for comparison. \**p*<0.05 (CircaCompare). Statistical significance reflects differences between baseline and post-shift within the same genotype.

In WT mice, the entrainment rate of *Per2* and *Bmal1* varied by tissue, with *Bmal1* in the liver realigning faster than *Per2*, while in WAT, both genes entrained at similar rates (WT; Δphase BL to D3, Liver; *Per2 p*<0.0001, *Bmal1 p*<0.0001, WAT; *Per2 p<*0.0001, *Bmal1 p<0.0001,* GLMMcosinor) (Fig. 2*B*, *C*, *F*). It is also worth noting that the rate of entrainment for the same clock gene varies across different tissues. WT *Per2* rhythms entrained more slowly in the liver than in WAT (WT; Δphase BL vs D5, liver *Per2* = 7.43 hours; WAT *Per2* = 5 hours) (Fig. 2*F*), while *Bmal1* entrained at comparable rates across both tissues (Fig. 2*B*-*E*). Irrespective of the differences in the rate of entrainment between clock gene rhythms within and between tissues, peripheral circadian oscillators in the liver and WAT were not entrained on D5 post shift in WT mice.

In contrast, *Per1^-/-^* mice displayed an accelerated entrainment of peripheral rhythms. By D3, *Per2* expression in the liver was fully re-aligned, with *Bmal1* exhibiting delayed entrainment, while in WAT, the pattern reversed (*Per1^-/-^*; Δphase BL to D3, liver; *Per2 p*=0.91 and *Bmal1 p*<0.0001; WAT; *Per2 p*<0.0001 and *Bmal1 p*=0.96; GLMMcosinor)(Fig. 2*F*).

Interestingly, unlike in the SCN where amplitude differences between WT and *Per1^-/-^* were not significant (Fig. S3, SCN *Bmal1* and *Per2*: D3 WT vs. D3 *Per1^-/-^*, *p*>0.05, D5 WT vs. D5 *Per1^-/-^*, *p*>0.05), peripheral oscillators in *Per1^-/-^* mice exhibited reduced amplitude during entrainment (Liver *Bmal1* and *Per2*: D3 WT vs. D3 *Per1^-/-^*, *p*<0.0005, D5 WT vs. D5 *Per1^-/-^*, *p*<0.0005; WAT *Bmal1* and *Per2*: D3 WT vs. D3 *Per1^-/-^*, *p*<0.0005, D5 WT vs. D5 *Per1^-/-^*, *p*<0.002; Fig. 2*B-E*). This oscillator instability, indicated by the proximity of *Per2* and *Bmal1* expression rhythms to the singularity point, may contribute to the observed acceleration in peripheral entrainment in *Per1^-/-^*mice.

Our results indicate that *Per1* modulates the entrainment dynamics of both central and peripheral circadian clocks, acting as a regulatory buffering mechanism for systemic physiological adaptation to environmental light shifts.

### *Period1* restricts the realignment rate of circadian rhythms in energy metabolism

Given the role of peripheral clocks in regulating daily metabolic rhythms ^7,30–32^, we hypothesized that the accelerated entrainment of circadian oscillators in the liver and WAT of *Per1*-deficient mice would result in faster realignment of systemic energy metabolism. To test this, we measured key systemic metabolic outputs—respiratory exchange ratio (RER), energy expenditure (EE), oxygen consumption (VO), and carbon dioxide production (VCO)—following a shifted light-dark (LD) cycle.

Following a 12-hour phase delay, *Per1^-/-^* mice exhibited significantly faster entrainment of metabolic rhythms compared to WT controls. Specifically, RER and EE rhythms in *Per1^-/-^*mice synchronized with the shifted LD cycle within two days, while WT mice required seven days to achieve a comparable alignment (PS_50_ WT vs. *Per1^-/-^*, RER; *p*=0.0003 and EE; *p*=0.0003, Student’s *t*-test) (Fig. 3*A,B,E* and *F*). Notably, the amplitude of RER and EE rhythms remained largely stable in *Per1^-/-^* mice during the entrainment process, with the exception of a transient decrease on D1 post-shift (Fig. 3*B,D* and *F*). By contrast, WT mice experienced a significant amplitude reduction during entrainment, which subsequently recovered once rhythms fully aligned with the new LD cycle (Fig. 3*B, F*). The reduction in RER rhythm amplitude in WT mice was primarily due to elevated trough levels (Fig. 3*C*), suggesting a shift towards increased carbohydrate utilization and reduced lipid oxidation during the entrainment period. These findings raise the intriguing possibility that dietary interventions targeting macronutrient utilization—such as modulating carbohydrate and lipid intake—could help mitigate jet lag by influencing metabolic rhythms and expediting alignment with new light-dark schedules.

**Fig. 3.**
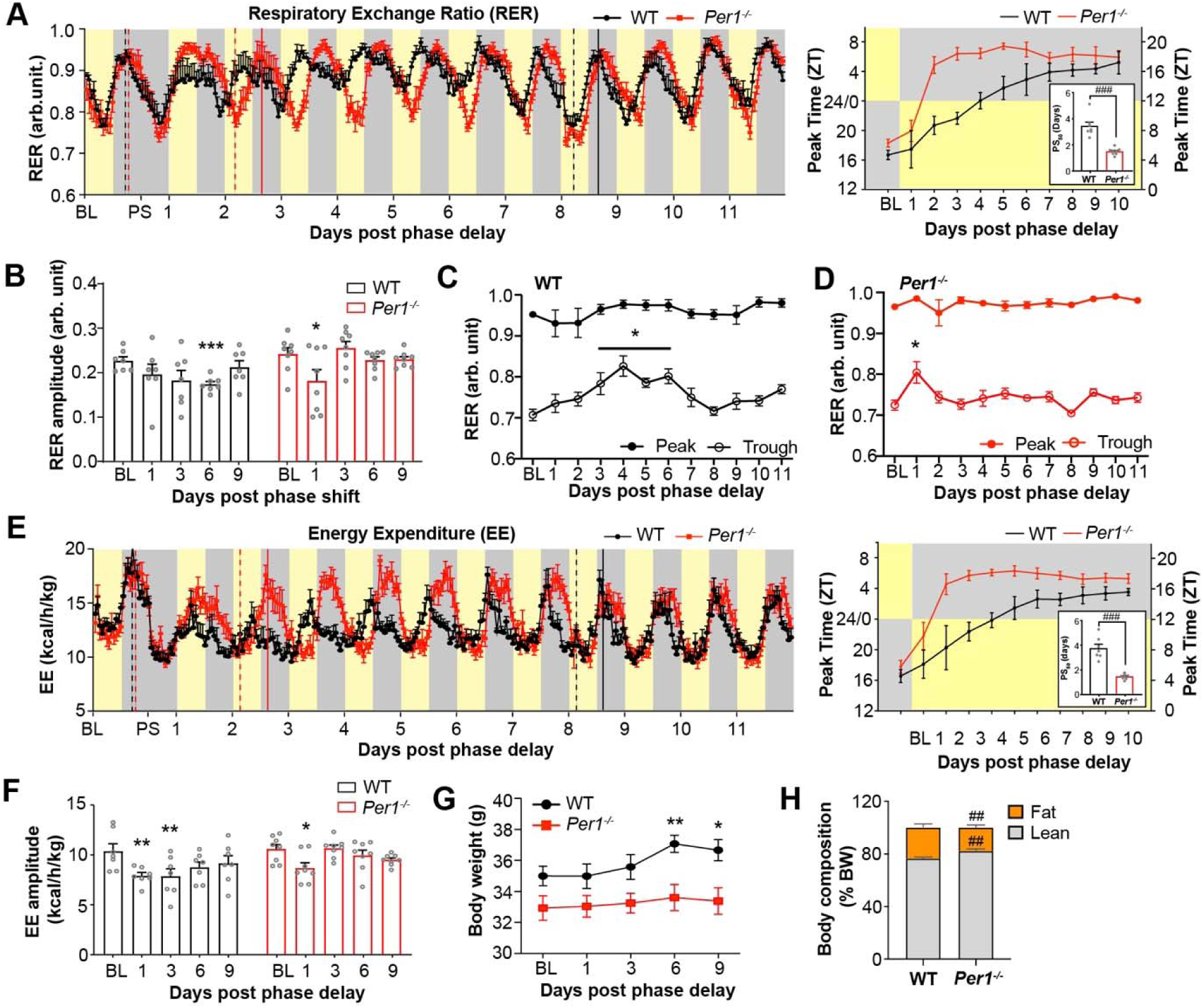
Altered energy metabolism response to jetlag in *Per1^-/-^* mice. **(A)** Respiratory exchange ratio (RER) rhythms in WT (black) and *Per1^-/-^*(red) mice before (BL) and after a 12-hour phase delay (PS). (Right) Average peak timing and PS_50_ for RER entrainment (p<0.0001, Student’s t-test). (**B**) Average RER rhythm amplitude from days 1–9 post-shift compared to BL in WT and *Per1^-/-^*mice (WT, *p*=0.031; *Per1^-/-^*, *p*=0.021; one-way ANOVA). (**C**–**D**) Peak and trough RER values before (BL) and post-shift (days 1–9) in (**C**) WT and (**D**) *Per1^-/-^* mice (WT, *p*=0.039; *Per1^-/-^*, *p*=0.043; two-way ANOVA). (**E**) Energy expenditure (EE) rhythms in WT and *Per1^-/-^*mice before and after the phase delay. (Right) Average peak timing and PS_50_ for EE entrainment (*p*<0.0001, Student’s *t*-test). (**F**) Mean EE rhythm amplitude before (BL) and after the phase delay in WT and *Per1^-/-^*mice (WT, *p*=0.0392; *Per1^-/-^*, *p*=0.0291; one-way ANOVA). (**G**) Body weight changes before (BL) and after the phase delay (WT, *p*=0.004; *Per1^-/-^*, *p*>0.05; one-way ANOVA). (**H**) Fat and lean body mass comparison post-shift in WT and *Per1^-/-^* mice (Fat, *p*=0.0043; Lean, *p*=0.0035; Student’s t-test). *n*=7 for WT, *n*=8 for *Per1^-/-^*. Data are presented as mean ± SEM. Statistical significance: \**p*<0.05, \*\**p*<0.005, \*\*\**p*<0.0005 (within genotype); #*p*<0.05, ##*p*<0.005, ###*p*<0.0005 (between genotypes).

Diurnal rhythms in VO and VCO levels in both genotypes were consistent with previous observations ^30^. While baseline characteristics of VO and VCO rhythms (amplitude, phase, and mesor) were similar between genotypes, *Per1^-/-^* mice displayed a significantly amplified response to the 12-hour LD phase delay. Specifically, VO and VCO rhythm amplitudes in *Per1^-/-^* mice were significantly elevated during the first five days (D5) of the LD shift (main effect of genotype, *p*=0.0013; two-way ANOVA followed by Sidak’s multiple comparisons between genotypes) (Fig. S4*A–D*). This amplitude increase in VO and VCO in *Per1^-/-^* mice was attributed to higher peak values, suggesting that *Per1* functions to buffer the metabolic oxidative response to environmental time shifts. The simultaneous increase in daily levels of VO_2_ and VCO_2_ also explains the increase in EE and RER rhythms amplitude during entrainment.

According to the principles of energy balance, weight gain reflects an imbalance between energy intake and energy expenditure ^31^. WT mice exhibited significant weight gain during the LD shift, coinciding with reduced RER and EE rhythm amplitudes (Fig. 3*H*). Importantly, this reduction in energy expenditure was not due to altered food intake; both WT and *Per1^-/-^* mice consumed similar amounts of food throughout the experiment (Fig. S4*E, F*). Unlike WT mice, *Per1^-/-^* mice maintained stable energy expenditure and body weight, with preserved body composition (WT: 76.53±1.17% of BW, *Per1^-/-^*: 82.21±0.44% of BW, *p*=0.0078, Two-way ANOVA) and lower fat accumulation upon full entrainment (WT: 23.48±1.17% of BW, *Per1^-/-^*: 17.81±0.44% of BW, *p*=0.0078, two-way ANOVA) (Fig. 3*G*, *H*). Notably, while daily food intake did not vary, WT mice displayed a disrupted feeding pattern, with intake shifted predominantly to the new light phase until D5 of entrainment (p=0.015; two-way ANOVA followed by Sidak’s multiple comparisons between the baseline and days after shift; Figs. S4*E,* F), whereas *Per1^-/-^* mice maintained their typical dark-phase feeding pattern throughout (main effect of time, *p*=0.38; two-way ANOVA; Figs. S4*E*, *F*).

These results indicate that *Per1* deletion mitigates jetlag-induced metabolic disruptions, including weight gain, by preserving stable feeding patterns, energy expenditure, and substrate utilization rhythms in alignment with the sleep-wake cycle. However, the molecular mechanisms driving this metabolic resilience remain unclear.

### *Period 1* shapes the diurnal rhythms of the liver metabolome

To determine whether this systemic metabolic stability extends to peripheral metabolic regulation, we focused on the liver, a key organ governing systemic energy homeostasis integrating circadian cues to coordinate energy balance. We hypothesized that *Per1^−/−^* mice maintain metabolic resilience through accelerated re-entrainment of central and peripheral clocks, enhanced clock plasticity, or a combination of both, which could stabilize clock-regulated metabolic pathways.

Building on evidence linking transcriptional and post-translational modifications to circadian disturbances ^32–34^, we postulated that *Per1* shields metabolic rhythms in the liver from external perturbations, much like its role in the SCN. To test this, we profiled the liver metabolome of *Per1^-/-^* and WT mice under stable (baseline) and 12-hour shifted (jetlag) LD cycles, collecting samples every 4 hours over 24 hours to capture diurnal variation in 276 metabolites entrained (baseline; BL) and jetlag conditions (D3 and D5 post-shift).

Under baseline conditions, 83.3% of metabolites (230 out of 276) were rhythmic in WT livers, while 94.2% (260 out of 276) showed rhythmicity in *Per1^-/-^*livers. The absence of *Per1* had minimal impact on the baseline metabolomic landscape, with 227 metabolites exhibiting conserved rhythmicity across genotypes (Fig. 4*A*-*C*). Focusing on pathways related to glucose and lipid metabolism, we observed that a larger proportion of bile acid metabolites were rhythmic in *Per1^-/-^*livers (197 of 209) compared to WT livers (167), along with rhythmic intermediates in the TCA cycle, short-chain fatty acids (SCFA), ketone bodies, and aldehydes. In WT livers, > 50 % of the rhythmic metabolite peaks were distributed across the 24-hour cycle, suggesting phase-coordination of metabolic pathways by the liver clock. Nevertheless, 41.7% of the rhythmic metabolites were phase-aligned with a peak at ZT20 (Fig. 4*E*). In contrast, 63.8% of rhythmic metabolites in *Per1^-/-^* livers peaked at ZT16, advanced by 4 hours compared to the WT peak at ZT20 (Fig. 4*E*). This shifted phase relationship between rhythmic key metabolic processes, such as bile acid synthesis, glycolysis (e.g., pyruvate), lipogenesis, ketogenesis (e.g., 3-hydroxybutyrate), and the TCA cycle, between genotypes, indicates that *Per1* shapes the timing of selective metabolic rhythms, ensuring their phase alignment under baseline conditions.

**Fig. 4.**
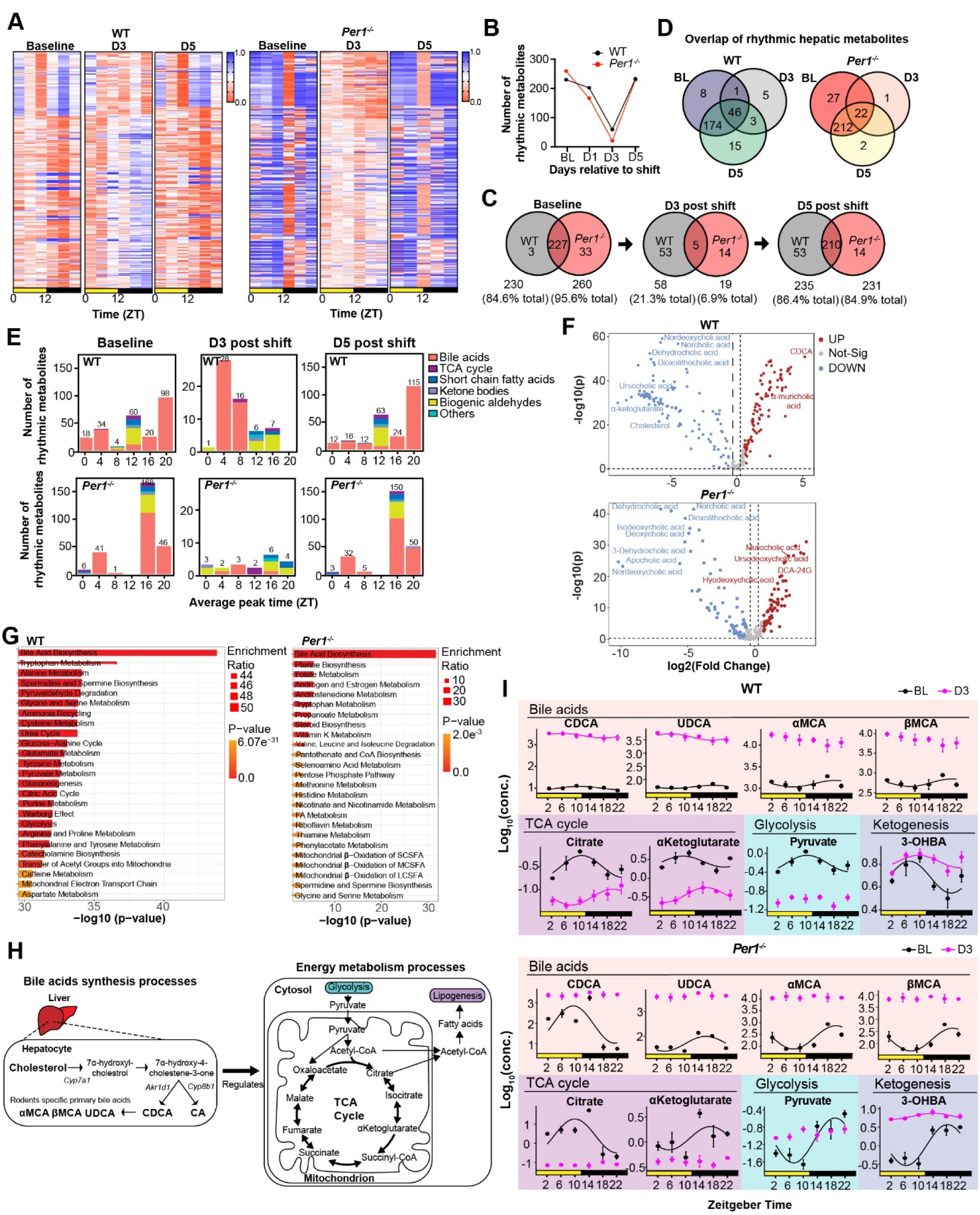
Impact of *Period1* on hepatic metabolome response to jetlag. (**A**) Heatmap of normalized hepatic metabolite expression in WT and *Per1^-/-^* mice at baseline (BL), day 3 (D3), and day 5 (D5) post 12-hour phase shift. (**B**) Total number of rhythmic metabolites at BL, D3, and D5 post-shift. (**C**) Venn diagrams showing metabolite changes at BL, D3, and D5 between genotypes. (**D**) Venn diagrams of rhythmic metabolites across BL, D3, and D5 within each genotype. (**E**) Peak phase distribution of rhythmic metabolites at BL, D3, and D5 in WT and *Per1^-/-^*mice. Numbers above bars indicate metabolite counts, color-coded by category. (**F**) Volcano plots of metabolites significantly altered (*p*< 0.05) by the phase shift in WT (top) and *Per1^-/-^* mice on D3 (bottom). (**G**) Enriched biological pathways of significantly altered metabolites post-shift, ranked by *p*-value. (**H**) Schematic of interconnected pathways involving selected metabolites. (**I**) Rhythms of selected metabolites at BL and D3 in WT and *Per1^-/-^* mice during re-entrainment. Data are presented as mean ± SEM. Statistical comparisons are between BL and post-shift days within each genotype using CircaCompare.

Our findings reveal that while *Per1* deletion minimally impacts the overall rhythmicity of liver metabolites, it significantly shifts the phase alignment of key metabolic pathways. This implies that *Per1* functions as a temporal regulator, coordinating the timing of liver metabolism to maintain phase coherence across interconnected metabolic processes. The observed phase shift in *Per1^−/−^* livers may reflect increased metabolic flexibility, potentially contributing to systemic resilience. Whether this altered metabolic timing affects the liver’s response to circadian misalignment remains unclear.

### The response of the liver metabolome to jetlag is strongly influenced by *Period 1*

Having established that *Per1* shapes rhythmic baseline (BL) liver metabolism, we next examined whether it stabilizes metabolic rhythms during circadian misalignment (Fig. 4). While *Per1*-deficient mice exhibited faster re-entrainment of systemic physiological rhythms—including the sleep-wake cycle, RER, and EE—the liver metabolome re-entrained at a similar rate in both genotypes. However, the sequence of realignment differed markedly.

In WT mice, the liver metabolome re-entrained before the alignment of liver clock gene expression, systemic metabolism, and the sleep-wake cycle. By contrast, in *Per1^−/−^* mice, metabolic re-entrainment only occurred after these broader physiological rhythms realigned (Fig. 5). These findings suggest that while the rate of liver metabolome re-entrainment is *Per1*-independent, *Per1* influences the temporal hierarchy of realignment across interconnected physiological systems.

**Fig. 5.**
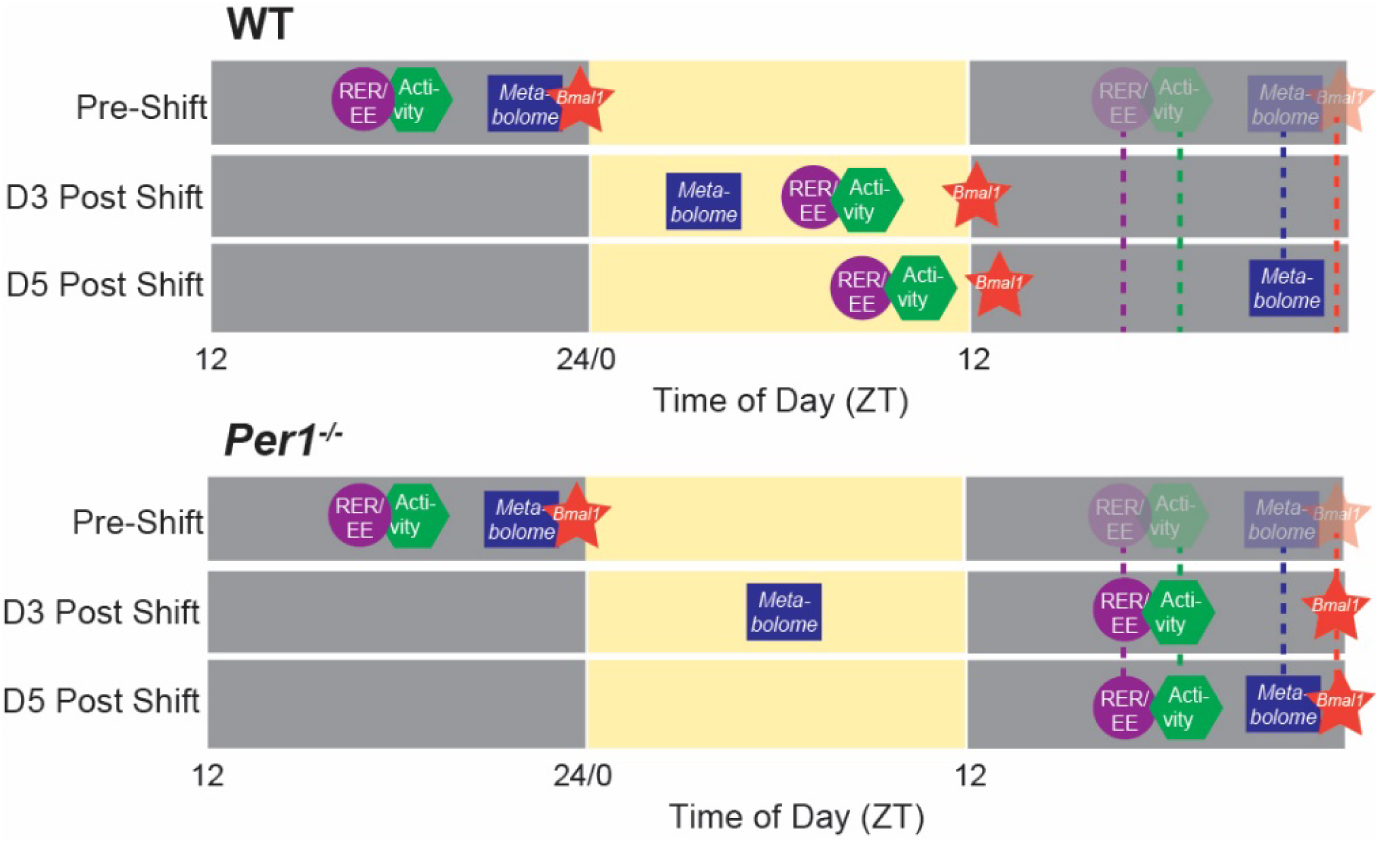
Schematic diagram showing entrainment rate of rhythms of activity, energy metabolism, *Bmal1* gene expression in the liver and its metabolome in WT and *Per1^-/-^* mice.

Closer inspection at the acute impact of jetlag on the liver metabolome revealed a substantial disruption of metabolite rhythmicity on D3 post-shift, with only 21.0% of metabolites retaining rhythmicity in WT mice, compared to 6.9% in *Per1^-/-^* mice (Fig. 4*C, D*). The rhythmic WT metabolites largely peaked around ZT4 on D3, suggesting gradual realignment to their baseline phase at ZT20 (Fig. 4*E*). In contrast, the remaining rhythmic metabolites in *Per1^-/-^*mice showed no phase coherence, peaking at various times throughout the day and night cycle. By D5, rhythmicity of most liver metabolites was restored in both genotypes (Fig. 4*A*-*C*), with phase alignment returning to baseline (Fig. 4*E*).

Jetlag also significantly affected absolute metabolite levels. On D3, nearly all metabolites were differentially regulated: 188 were upregulated and 240 downregulated in WT mice, while 176 were upregulated and 246 downregulated in *Per1^-/-^* mice (Fig. 4*F*). Pathways critical for energy metabolism, such as glycolysis, amino acid metabolism, fatty acid oxidation, glucose metabolism, the TCA cycle, and bile acid synthesis, were markedly impacted in both genotypes (Fig. 4*G*). Key metabolites like chenodeoxycholic acid (CDCA), ursodeoxycholic acid (UDCA), α-muricholic acid (αMCA), β-muricholic acid, citrate, α-ketoglutarate, and pyruvate exhibited altered rhythmicity on D3 (Fig. 4*H*). Notably, in *Per1^-/-^*mice, most metabolites, except for 3-hydroxybutyrate (3-OHBA), became arrhythmic on D3, whereas WT mice retained rhythmicity in several of these critical metabolites. This finding indicates that *Per1* stabilizes specific metabolic pathways during circadian re-entrainment.

The pronounced metabolic instability in *Per1^−/−^* mice proposes that *Per1* buffers metabolic rhythms against environmental time shifts. While feeding rhythms are key zeitgebers for liver metabolism, the rapid re-entrainment of sleep-wake and feeding cycles in *Per1^−/−^* mice by D3 suggests that additional entraining mechanisms regulate liver metabolome rhythmicity in the absence of *Per1*.

These findings suggest that *Per1* strengthens the coupling of the liver metabolome to the local hepatic clock, ensuring metabolic pathways remain rhythmic under shifting light-dark schedules. In its absence, metabolic rhythms become more dependent on systemic cues, leading to increased phase instability despite the rapid re-entrainment of systemic metabolic and clock gene rhythms. Together, our findings establish *Per1* as a critical regulator of hepatic metabolic rhythms, maintaining phase coherence under stable conditions and buffering against circadian misalignment.

This pronounced instability of the liver metabolome underscores the fundamental differences in how the SCN and peripheral clocks respond to circadian misalignment. In contrast to the liver, where timekeeping is primarily governed by autonomous molecular oscillations with relatively weak intercellular coupling, the SCN functions as a highly interconnected network of coupled oscillators, enabling robust phase coordination and resilience to perturbations^35–38^. To investigate whether *Per1* plays a role in this SCN-specific coupling and contributes to its entrainment dynamics, we next examined the SCN’s responsiveness to light, the primary zeitgeber regulating circadian adaptation.

### *Period1* modulates light-induced activation of SCN neurons

To investigate the mechanisms underlying accelerated circadian realignment in *Per1^-/-^* mice, we first examined whether *Per1* influences photoreception. The retina, which contains intrinsically photosensitive retinal ganglion cells (ipRGCs) expressing melanopsin (OPN4), transmits photic signals to the SCN to synchronize circadian rhythms^39^. We quantified melanopsin-positive (OPN4) ipRGCs in flat-mounted retinas and found no significant difference between genotypes (WT vs. *Per1^-/-^ p*=0.8581, Student’s t-test) (Fig. 6*A, B*), indicating that genotype-specific differences in photoentrainment are not due to variations in ipRGC abundance.

**Fig. 6.**
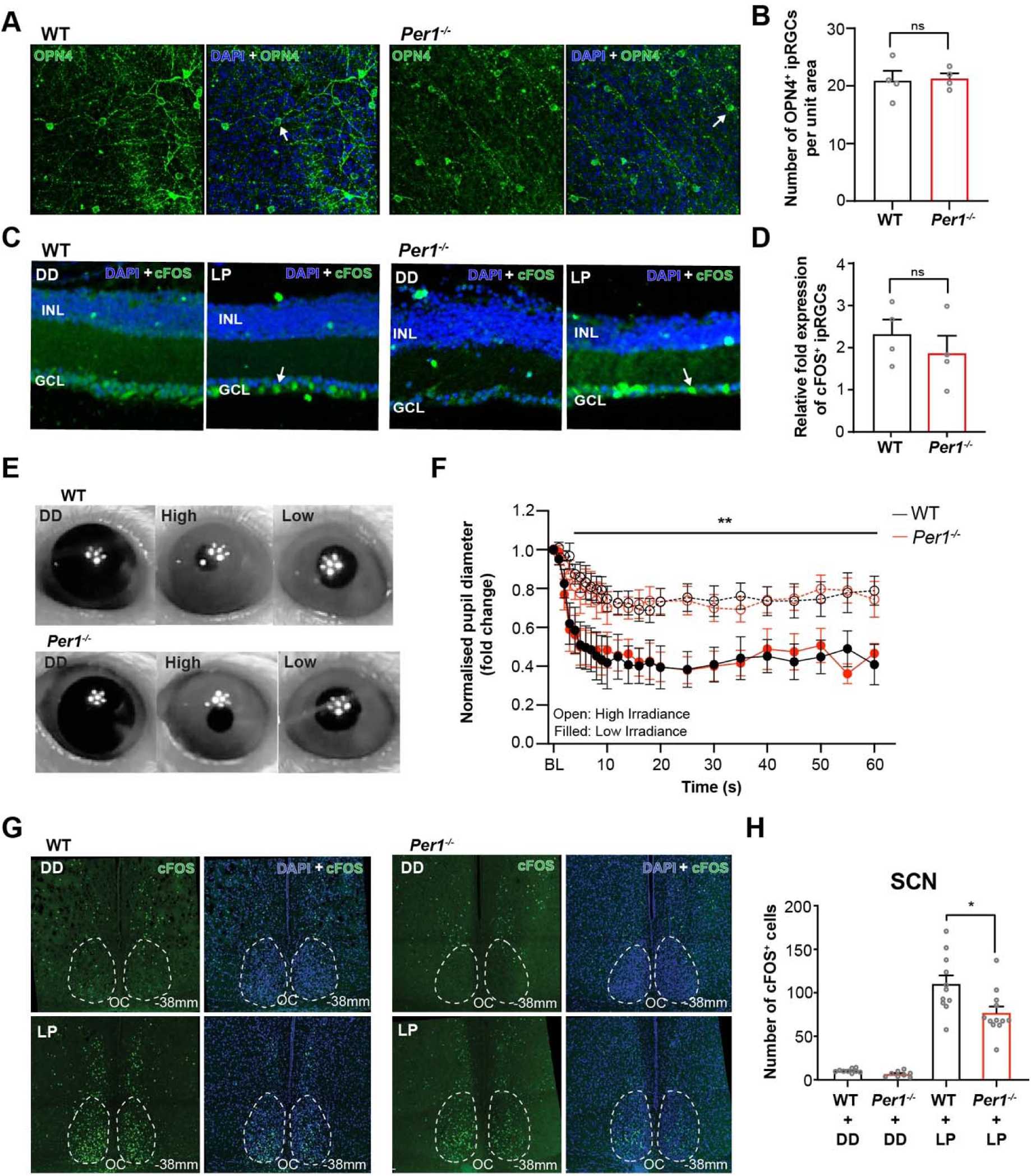
*Period1* does not alter retinal photosensitivity to light. (**A**) Confocal images of melanopsin (OPN4) expression in intrinsically photosensitive retinal ganglion cells (ipRGCs) in medial retina sections of WT (left) and *Per1^-/-^* (right) mice. (**B**) OPN4^+ve^ ipRGC cell counts per 0.14 mm² in medial and lateral retina sections of WT (black) and *Per1^-/-^* (red) mice. (**C**) Confocal images of cFOS expression in the retinal ganglion cell layer following a 1-hour light pulse at CT14 in WT (left) and *Per1^-/-^* (right) mice. (**D**) Quantification of cFos signal in cFOS^+ve^ ipRGCs in response to light pulse relative to the control (dark conditions). (**E**) Pupillary light reflex in WT and *Per1^-/-^* mice under high and low irradiance. (**F**) Pupil constriction (%) in response to bright light, relative to dark-adapted pupil diameter (High vs. Low irradiance, *p*<0.005, two-way ANOVA; *n*=6 WT, *n*=5 *Per1^-/-^*). (**G**–**H**) Confocal images and quantification of cFOS in the SCN under dark and light pulse conditions (LP; WT vs. *Per1^-/-^*, *p*=0.012, Student’s *t*-test; *n*=4 for each genotype). Data are presented as mean ± SEM. Statistical significance: \**p*<0.05, \*\**p*<0.005, ns *p*>0.05.

We then assessed the functional response of the retinal ganglion cell layer by measuring cFOS induction, a marker of light-activated neural activity, following a 1-hour light pulse at 35 lux administered at CT14. Both WT and *Per1^-/-^* mice displayed significant cFOS induction (cFOS induction: WT, *p*=0.0191 and *Per1^-/-^*, *p*=0.0419) (Fig. 6*C*), with no significant differences in cFOS cell counts between genotypes (WT vs. *Per1^-/-^ p*=0.4345) (Fig. 6*D*). Additionally, the pupillary light reflex (PLR) was measured as a functional indicator of ipRGC activation. Both genotypes showed significant pupillary constriction in response to bright (250 lux) and dim (60 lux) light, with no genotype-dependent differences (*p*<0.0001, two-way ANOVA) (Fig. 6*E, F*). These findings suggest that *Per1* does not modulate retinal responsiveness to light.

We next examined SCN neuron activation by measuring cFOS expression in response to a 1-hour light pulse at CT14. While light significantly induced cFOS in SCN neurons of both genotypes (DD vs. LP, WT *p*<0.0001; *Per1^-/-^ p*<0.00001, Student’s t-test) (Fig. 6*G*), *Per1^-/-^* mice showed a 30% reduction in cFOS induction compared to WT (*p* = 0.015) (Fig. 6*H*). The negative correlation between cFOS induction and accelerated behavioral and metabolic re-entrainment measures in *Per1^-/-^* mice was unexpected. Based on our findings, cFOS induction does not integrate photic information nor predict behavioral phase shifts. Instead, it may be part of the *Per1*-dependent buffering mechanism against excessive light-induced SCN activation.

### *Period1* regulates intercellular coherence in the SCN

The diminished light responsiveness of SCN neurons in *Per1*^-/-^ mice indicates that *Per1* influences SCN network integrity, potentially through effects on molecular oscillator stability or intercellular synchrony. To investigate this, we examined *Per1*’s role in maintaining intercellular coherence within the SCN using spatial transcriptomics to identify gene expression differences between WT and *Per1*^-/-^ mice (Fig. 7*A*). Among the over 9,000 detected genes, 11 showed significant differential expression (Fig. 7*B–D*). Notably, *Avp* (arginine vasopressin) and *Pcsk1n* (proprotein convertase subtilisin/kexin type 1 inhibitor)—both linked to SCN interneuronal coupling—were down-regulated in *Per1^-/-^* mice (Fig. 7*D, E*). AVP is essential for SCN intercellular coherence ^40,41^, while *Pcsk1n* modulates neuropeptide signaling within SCN core neurons ^42^. In situ validation using RNAscope confirmed reduced *Avp* expression in *Per1^-/-^* SCN but found no significant difference in *Pcsk1n* (Fig. 7*F, G* and Fig. S5). AVP protein levels were also significantly lower in *Per1^-/-^* mice compared to WT mice (*p*<0.0005; Fig. 7*H–J*), supporting the hypothesis that *Per1* modulates SCN intercellular coupling, thereby impacting circadian entrainment. These findings provide a mechanistic basis for *Per1*’s role as a temporal buffer, linking AVP-dependent SCN coupling to the broader systemic regulation of circadian rhythms.

**Fig. 7.**
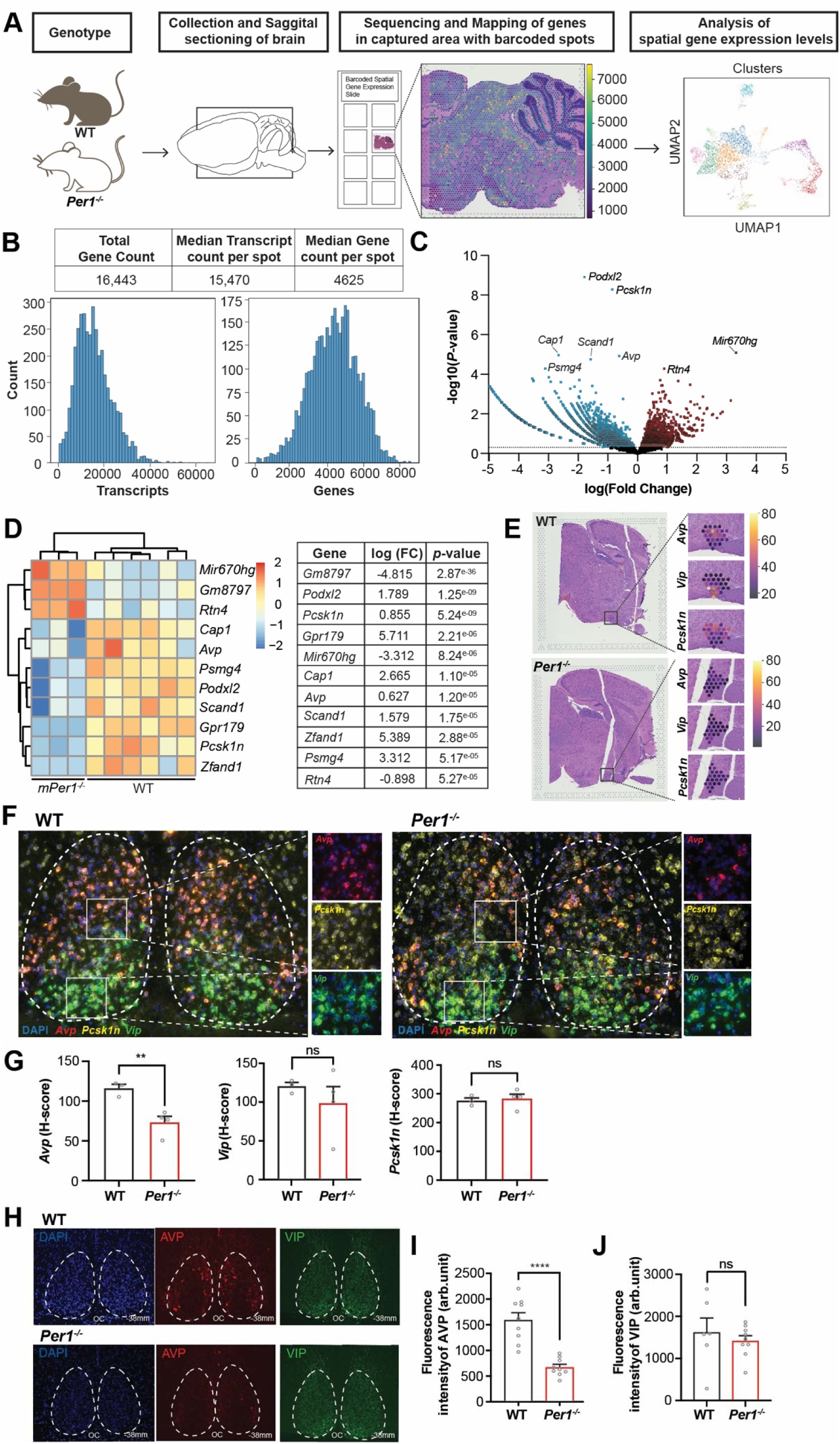
Loss of *Period1* alters transcriptomic profiles in SCN neurons. (**A**) Workflow for spatial transcriptomics from tissue collection to data analysis. (**B**) Total and median gene counts in sagittal brain slices of WT and *Per1^-/-^*mice. (**C**) Volcano plot showing significantly upregulated (red) and downregulated (blue) genes in the SCN of *Per1^-/-^* mice. Top differentially expressed genes are labeled. (**D**) (Left) Heatmap of the top 11 significantly altered genes in WT and *Per1^-/-^* mice. (Right) Table of fold changes and p-values. (**E**) Spatial expression profiles of genes related to SCN interneuronal synchrony (*Vip*, *Avp*, *Pcsk1n*). (**F**) RNAscope images showing RNA signals for *Vip*, *Avp*, and *Pcsk1n* in SCN slices of WT and *Per1^-/-^* mice. (**G**) Quantification of RNA counts for *Vip*, *Avp*, and *Pcsk1n* (*Avp*, *p*=0.0084, Student’s *t*-test; *n*=4 WT, *n*=3 *Per1^-/-^*). (**H**) Immunofluorescence images of VIP and AVP proteins in SCN slices from WT and *Per1^-/-^* mice. (**I**–**J**) Quantification of fluorescence intensity for VIP and AVP (AVP, *p*<0.0001, Student’s *t*-test). Data are shown as mean ± SEM. Statistical significance: \*\**p*<0.005, \*\*\*\**p*<0.0001.

Taken together, our results demonstrate that *Per1* functions as a gatekeeper of circadian stability, buffering the circadian system against over-responsiveness to environmental light cues. By orchestrating central SCN coupling, peripheral clock alignment, and systemic metabolic rhythms, *Per1* emerges as a critical modulator of circadian adaptation with significant implications for mitigating jetlag and circadian misalignment.

## Discussion

Circadian clocks must balance adaptability with stability—adjusting to environmental changes without overreacting to noise. Our findings identify *Per1* as a key regulator of this balance. Rather than facilitating re-entrainment, *Per1* acts as a molecular brake, slowing phase shifts to preserve synchrony across brain and body. In its absence, re-entrainment is accelerated but less coordinated, revealing a fundamental trade-off between speed and stability in circadian adaptation.

A key aspect of circadian photoentrainment is the controlled, gradual adjustment of internal rhythms to external light cues, a process essential for preventing abrupt phase shifts in response to environmental changes ^8^. These inherent limitations explain why we experience jetlag or struggle with shift work, as the circadian system prioritizes robustness over plasticity. This balance, favoring stability to avoid overreacting to subtle changes, comes at the cost of slower adaptation to necessary shifts, such as light-induced phase shifts. In both humans and mice, this robustness constrains the rate of entrainment. Although the exact mechanism regulating this balance remains unclear, our results suggest that PER1 plays a crucial role in this process, offering a potential intervention to enhance circadian plasticity. Targeting PER1 function may improve the efficacy of phototherapy for jetlag, circadian disruption in neurodegenerative diseases ^43^, and metabolic disorders^44–46^.

The gene *Per1* has long been identified as crucial in resetting the circadian clock in response to light. *Per1* functions as an immediate early gene, induced by nighttime light exposure ^24^, which shifts the SCN clock and, in turn, adjusts downstream rhythms, including the sleep-wake cycle. The expression of *Per1* correlates with the magnitude of these light-induced phase shifts in the circadian clock, and its induction varies depending on the time of day ^22–24^.

Several studies suggest that the circadian clock’s sensitivity to light can be enhanced through pharmacological manipulation. For example, inhibiting SIK1—both in vitro and in vivo—has been shown to increase *Per1* expression in response to phase-shifting stimuli, suggesting that SIK1 acts as a repressor. By dampening *Per1* induction, SIK1 may help regulate how the circadian clock responds to phase-shifting stimuli ^47^. Similarly, mice deficient in *Id2* exhibit faster adjustment to changes in light schedules, a phenomenon accompanied by increased light-induced *Per1* expression in the SCN (Duffield et al., 2009; Duffield et al., 2020). Additionally, Akiyama et al. (1999) found that blocking *Per1* expression with antisense oligonucleotides prevented behavioral phase delays in response to nighttime light pulses.

Collectively, these studies suggest that *Per1* induction is integral to how the molecular clock perceives and responds to light. However, misconceptions persist about the necessity of *Per1* for photoentrainment. Pendergast and Yamazaki demonstrated that *Per1* is not required for light-induced behavioral phase-shifts; in their study, *Per1*-deficient mice still responded to phase-shifting light pulses. They speculated that the absence of *Per1* could even accelerate entrainment to shifted light-dark schedules, implying that other pathways might compensate for *Per1* loss ^48^.

In this study, we test this hypothesis directly, providing a detailed examination of *Per1*’s role in light entrainment. Consistent with Pendergast et al. (2010)^48^, we found that *Per1^-/-^* mice exhibit both phase-delaying and phase-advancing responses to light pulses. Notably, *Per1^-/-^* mice displayed phase delays nearly four times greater than wild-type (WT) mice after a light pulse at CT14, while the phase advance response at CT19 was similar between genotypes.

Interestingly, our findings differ from those of Albrecht et al. (2001) ^22^, who reported that a different *Per1* knockout strain (*Per1^Brdm1^*) showed phase delays similar to WT mice at ZT14. These differences may be due to variations in the genetic modifications used to disrupt *Per1* function. In *Per1^Brdm1^* mice, exons 4 through 18 were replaced with the *Hprt* transgene, producing a mutated transcript likely rendered non-functional by a disrupted N-terminal sequence ^49^. Our study utilized *Per1^Idc^* transgenic mice, in which exons 2 through 12 were replaced with a neomycin cassette, resulting in a non-functional *Per1* product ^16^.

Amplified behavioral shifts in locomotor activity rhythms in response to a phase-delaying light pulse led us to question whether such a response translates into accelerated adaptations of *Per1*-deficient mice to changes in the photo-schedule. Indeed, we found that *Per1^-/-^* mice re-entrain to a 12-hour phase delay quickly and reach full entrainment of the sleep-wake cycle within two days. This behavioral phenotype is not due to a masking effect by light, but due to immediate entrainment of the SCN clock, as evidenced by the rapid phase re-alignment of core clock gene expression rhythms in *Per1^-/-^*mice.

Our findings demonstrate that *Per1* deletion accelerates circadian realignment but compromises SCN network coherence. Under a reversed light-dark (LD) schedule, *Per1^-/-^* mice rapidly aligned their sleep-wake rhythms, paralleling faster entrainment of the SCN molecular oscillator. However, when released into constant darkness (DD) after just one reversed LD cycle, *Per1^-/-^* mice exhibited arrhythmicity in their sleep-wake behavior (Fig. S6), unlike the stable free-running rhythms observed in WT mice. This instability was mirrored at the molecular level, where *Bmal1* expression in the SCN of *Per1^-/-^* mice briefly lost rhythmicity on the first day of LD reversal before recovering and entraining to the shifted cycle by day three (Fig. S3). In contrast, WT SCN rhythms transitioned gradually, reflecting stronger intercellular coupling and a slower but more stable adaptation process.

These observations highlight a trade-off: *Per1* loss enhances circadian flexibility, enabling rapid phase shifts, but at the cost of transient instability. By maintaining intercellular synchrony, *Per1* buffers the SCN clock against excessive light-induced resetting, stabilizing rhythms in the face of environmental perturbations. This dual role reveals how the circadian system balances robustness with adaptability—a critical mechanism for responding to sudden changes while preserving biological stability.

To uncover the mechanism underlying *Per1*’s buffering effect, we investigated its role in SCN synchrony using spatial transcriptomics. We identified significant downregulation of *Avp* in *Per1^-/-^*mice, a neuropeptide essential for intercellular coupling in the SCN ^40^ through autocrine signaling via V1a and V1b receptors ^50^. Prior studies have shown that AVP-deficient mice exhibit rapid re-entrainment to shifted LD cycles ^41^, mirroring the enhanced circadian plasticity seen in *Per1^-/-^* mice. This suggests that reduced *Avp* expression weakens intercellular communication, compromising SCN stability and rendering the central clock more sensitive to environmental light cues.

The link between *Per1* and AVP provides a molecular explanation for the accelerated yet unstable circadian shifts in *Per1^-/-^*mice. While reduced intercellular coupling allows for faster adaptation, it also introduces transient desynchrony—a trade-off between adaptability and robustness. These findings position *Per1* as a critical stabilizer of SCN network function, where its regulation of AVP and intercellular coupling buffers the circadian system against excessive light-induced perturbations, ensuring rhythm resilience amidst environmental fluctuations.

The molecular mechanism of photoentrainment involves transcriptional activation of downstream target genes including *Per1, cFos*, *Nr4a1* and many more, mediated by cAMP response element binding protein (CREB) phosphorylation ^51–54^. Therefore, immediate induction of *Per1* and *cFos* by light is often linked to light-regulated clock resetting ^55–57^. Here, we observed significantly attenuated induction of cFOS in *Per1^-/-^* SCNs by light. Given the critical role of AVP in interneuronal communication and SCN oscillator coupling, it is reasonable to speculate that low expression of AVP in *Per1^-/-^* SCN creates a more chaotic state within the SCN, with reduced internal synchrony. The resulting phase shifts in individual SCN oscillators could lead to variable responses to a stimulus. Indeed, with a CT14 light pulse, the induction of SCN cFOS expression is significantly reduced in *Per1*-deficient mice compared to WT. When profiling the spatial distribution of CREB activation (pCREB), an organized spatial map emerges in the pCREB signal in WT SCNs compared to the more random profile observed in SCNs of *Per1^-/-^*-mice (Fig. S7). This could be attributed to either reduced interneuronal coupling in the SCN or the autoregulatory feedback mechanism of *Per1*, which we previously discovered ^53^.

In *Per1*^-/-^ mice, clock gene expression rhythms in the SCN and peripheral organs, aligned within five days post-shift, correlated with stable energy metabolism rhythms. Notably, rapid re-entrainment to phase delay protected *Per1^-/-^* mice from the metabolic disturbances and weight gain observed in WT mice. In WT mice, SCN rhythms aligned faster than peripheral tissues (e.g., liver and fat), causing temporary circadian network disruptions and metabolic instability, including altered lipid oxidation, energy expenditure, and weight gain. These metabolic disturbances in WT mice likely stem from misalignment between feeding rhythms and internal clocks during entrainment ^58–60^. Thus, targeting *Per1* signaling provides physiological and metabolic resilience under jetlag conditions. These results collectively reveal that the silencing of *Per1* signaling, which has no phenotype under entrained conditions and upon acute LD cycle reversal, generates significant physiological advantages and becomes metabolically protective under jetlag conditions.

Overall, although *PER1* itself may not be readily druggable, its associated regulatory pathway, including AVP signaling and CREB-mediated transcription, offers feasible points of intervention for future chronotherapeutic strategies.

The liver is crucial in regulating the rhythmic manifestation of metabolic pathways, mainly controlling the synthesis and metabolism of glucose, lipids, and bile acid ^44^. Using metabolomics, we found that the liver metabolome in *Per1*^-/-^ mice decoupled from the liver clock during re-entrainment, showing increased arrhythmicity under jetlag conditions. Interestingly, in *Per1^-/-^* mice, the re-entrainment of the liver metabolites occurs after the sleep-wake cycle re-aligns, which may suggest that peripheral circadian rhythms are listening more closely to behavioral inputs like the sleep and wake signals in the absence of *Per1*.^61,62^.

Our study is unique in mapping dynamic changes within the circadian network that spans the brain and peripheral tissues, including the liver, adrenal, and fat. By analyzing specific nodes within this network, we provide a comprehensive view of *Per1*’s role in synchronizing both central and peripheral clocks and coordinating metabolic outputs. This approach allowed us to capture the systemic effects of *Per1* deletion on whole-body physiology, offering a blueprint for evaluating circadian clock gene manipulation as a therapeutic strategy.

Evidence suggests that the role of *Per1* is evolutionarily conserved, with the gene subject to stringent selection pressures that underscore its functional importance—likely explaining why mutations in *hPer1* are so rare. This evolutionary constraint may reflect *Per1*’s protective role in maintaining robust circadian rhythms, as demonstrated in our findings. Notably, the rare *hPer1* T2434C polymorphism has been linked to extreme diurnal preference, connecting *hPer1* to behavioral responses to light ^63^. In humans, diurnal preference (or chronotype) strongly influences an individual’s response to the phase-shifting effects of light, as shown in studies on abrupt time zone changes and shift (night) work ^64,65^. These findings support a role for *Per1* in mediating the clock-resetting effects of light in humans.

Modern society’s growing reliance on rotational shift work to deliver more convenient and efficient services has led to a surge in circadian rhythm and sleep disturbances ^66,67^. The chronic misalignment between an individual’s sleep-wake cycle and their internal biological processes has been strongly linked to a variety of health issues, with a particularly high risk of metabolic diseases. There is compelling evidence that shift work is a significant factor contributing to the rising prevalence of conditions like type 2 diabetes ^68–70^. In light of our new research findings, the *Per1* gene and PER1 protein emerges as promising therapeutic targets that could potentially mitigate the adverse effects of prolonged internal desynchrony, reducing the risk of shift-work-related diseases and easing the burden on healthcare systems.

## Materials and Methods

### Animal housing

Mice with deletion of *Per1* (*Per1^-/-^*) were obtained by crossing *Per1^-/-^* mice generated by with C3H/HeN mice and bred for at least 10 generations. Wild-type (WT) C3H/HeN littermates were used as controls. Male WT and *Per1^-/-^* mice (3 – 4 months of age) were individually housed in constant conditions of temperature (25 C°) and humidity (50%) with *ad libitum* access to water and standard chow. Before exposure to experimental conditions, mice were kept under a 12-hour light:12-hour dark cycle (LD 12:12) with white light (250 lux) in circadian cabinets (Phenome Technologies Inc., IL, USA) unless otherwise stated. Under the LD 12:12 condition, *Zeitgeber* time (ZT) was used to indicate light and dark onset - ZT0 and ZT12 correspond to light and dark onset, respectively.

### Locomotor activity monitoring

To determine the rate of photoentrainment, we used activity rhythms acquired via two methods: running wheel activity and homecage activity. Running wheel data of mice housed in running-wheel fitted cages was captured using the software Clocklab (ActiMetrics, IL, USA). Horizontal locomotor activity was captured using infrared cameras (Rasberry Pi NoIR Camera Module v2, RS Components Pty Limited, NSW, Australia). The video data was collected at a resolution of 640 × 480 pixels at 30 frames per second, and an automated frame-by-frame comparison was performed every 200 ms to determine movement (≥1 % change between frames was registered as movement).

### Measuring behavioral entrainment

#### For light pulse experiments

To examine light-induced clock resetting, WT and *Per1^-/-^*mice maintained under LD 12:12 were left in constant darkness (DD) starting from the dark phase of the day before the exposure to a light pulse, which is the baseline day. Twenty-four hours later, both genotypes received a single 1-hour light pulse (35 lux to minimize negative masking) at CT14 or CT19. Mice continued in DD for at least one additional week. The magnitude of the activity-onset rhythm phase shift induced by the light pulse was measured for each genotype.

#### For jet-lag experiments

To assess the rate of behavioral entrainment, WT and *Per1^-/-^*mice were subjected to a 12-hour shifted LD cycle. To reverse the LD cycle, the original nighttime was extended by 12 hours. The next 24-hour starting with the new light-onset of the reversed 12:12h LD cycle was defined as Day 1 (D1) (diagram in Figure 1A). The illumination for the 12-hour light phase was set to 35 lux (top of the cage) to minimize negative masking. The 50% phase shift (PS_50_) was determined to measure the rate of adaptation to the reversed light schedule by fitting a nonlinear regression line to the locomotor activity-onsets. The length of time required to reach the midpoint of circadian alignment is the PS_50_.

### Pupillometry

Consensual pupillary light reflex was measured to investigate the role of *Per1* in the function of the melanopsin-regulated nonimage-forming visual pathway. Mice were dark-adapted for 1-hour before light exposure at ZT8 using high irradiance (250 lux) and low irradiance (60 lux). Animals were placed so that the eye subjected to light was facing the recording infrared camera (Raspberry Pi NoIR night camera), capturing images at 30 frames per second for 1 minute. The video images of the eye were acquired at multiple time points over a minute to measure pupil constriction using Adobe Media Encoder 2022 (Adobe Creative Cloud, CA, USA). The pupil constriction was quantified by measuring the relative pupil size during light exposure normalized to the dilated pupil size in darkness, as previously described.

### Electroencephalography (EEG) and sleep data acquisition

Mice were deeply anesthetized with isoflurane (4% for induction and 2% for maintenance during the procedure). An incision was made from the head to midway between the scapulae. The EEG recording electrode was implanted 1.0mm anterior and 1.0mm lateral to the Bregma and the reference electrode was positioned 1.5mm posterior and 1.5 lateral to the Bregma on the contralateral side. The electromyography (EMG) recording electrodes were implanted in the trapezius muscle. The transmitter was placed subcutaneously along the dorsal flank between the forelimb and the hind limb. Upon recovery from implantation, mice were housed in standard cages and placed on a receiver to acquire data through the Ponemah data acquisition system. The transmitters, receivers, and Ponemah were purchased from Data Science International (DSI, MN, USA). The acquired data were processed using NeuroScore (DSI, MN, USA) to analyze vigilance states quantitively and qualitatively (i.e., Wake, NREM, and REM sleep).

### Energy metabolism

Mice were individually housed in climate-controlled automated metabolic cages (Phenomaster, TSE Systems, Germany) equipped with infrared sensors to capture horizontal locomotor activity, to monitor food and fluid intake, and for indirect calorimetry measurements (including respiratory exchange ratio (RER), energy expenditure, body weight change, oxygen consumption (VO_2_), and carbon dioxide production (VCO_2_). The mice were acclimatized to the chambers for at least one week prior to exposure to the 12-hour phase delay and remained in the chambers until the termination of experiments. Pre- and post-shift data were continuously acquired for the duration of the experiment (until mice were fully entrained to the new light schedule).

### Profiling rhythms in clock gene transcripts

To assess the baseline rhythm of clock genes, mice maintained under LD 12:12 were sacrificed, starting from ZT2, every 4-hours over a 24-hour period. To evaluate the entrainment dynamics of clock gene rhythms, mice that were phase shifted by 12 hours (as described above) were sacrificed on days 1, 3 and 5 (D1, D3 and D5) post-shift every 4 hours over a period of 24 hours starting from ZT2 of the new lighting schedule. Fresh brains, liver, and white adipose tissue (WAT) were collected and frozen until further processed. The SCN region from fresh-frozen brains was extracted in ice-cold RNAlater (Thermo Fisher Scientific, #AM7020) and stored at −80°C. The tissues were homogenized, and the RNA was extracted using RNAeasy micro kit (Qiagen, #74004). The cDNA was synthesized using Invitrogen High-Capacity cDNA reverse Transcription Kit (ThermoFisher Scientific, #LTS4368813). qPCR was performed on a Quantstudio7 (Applied Biosystems). The expression levels of the genes of interest were quantified using Taqman Gene Expression Assays and Taqman Master Mix (Applied Biosystems, #LTS4369016). The genes of interest were *Bmal1* (Mm00500226_m1) and *Per2* (Mm00478099_m1), and *Tbp* (Mm01277045_m1) was used as a housekeeping gene. All kits and assays were performed according to the manufacturer’s instructions.

### Immunohistochemical analysis of light-induced activation of the retina and SCN

After exposing mice to a single 1-hour light pulse at CT14 (35 Lux), mice were transcardially perfused. At the time of perfusion, the mice were deeply anesthetized and perfused with 0.9 % saline, followed by fixation in 4 % paraformaldehyde (PFA) in phosphate buffered saline (PBS). Fixed brains and eyes were collected and cryoprotected in 15 % sucrose and then 30 % sucrose solution in PBS at 4°C overnight. The cryoprotected brains and eyes were then frozen and cryo-sectioned at a 16 μm. Some of the cryoprotected retinas were flat-mounted for further processing. For immunostaining, the tissues were washed in PBS and permeabilized for antigen retrieval by heating the tissue in citrate buffer (pH 6) in a microwave. The tissue slides were blocked in bovine serum albumin (BSA) diluted in PBS with 0.3 % Triton X-100 (PBST) (0.02 g BSA/mL PBST) for an hour at room temperature. After blocking the tissue, the slides were incubated in blocking solution containing primary antibodies at 4°C overnight. The primary antibodies used were rabbit anti-cFOS antibody (1:1,500, Santa Cruz Biotechnology, #sc-166940) and rabbit anti-OPN4 antibody (1:2,000, Advanced Targeting Systems, AB-N38). The following day, slides were washed 10 minutes in PBST three times and incubated in secondary antibody, Alexa 488-conjugated goat anti-rabbit antibody (1:1,000, Invitrogen, #A-11078). Brain slices were washed as above and VectaShield with DAPI (Vector Laboratories, H-1200) was applied to visualize nuclei and stored at 4°C until confocal imaging.

### Image acquisition and cell counting

The regions of interest (ROIs) were imaged using a Leica DMi8 SP8 inverted confocal microscope and Leica Application Suite X software. A 20x objective was used to capture 1024×1024 images of all ROIs (average 15 stacks per image with a z-stack size of 1.75μm). Laser exposure, detector gain, pinhole, and offset were kept consistent across all samples. The acquired images were further processed using Imaris 8.4.1 software (Bitplane) in a 3-dimensional space. ROIs in brain sections were outlined using the top 5 brightest consecutive layers from z-stack images. Background noise was removed using the automatic threshold background subtraction method on Imaris. Only cells that met the following criteria were counted. For both cFOS and OPN positive cells in the SCN and retina, respectively, the intensity of the fluorescent signal had to be above the automatic threshold determined by Imaris and colocalize with DAPI-stained nuclei. The total number of cFOS or OPN-positive cells was normalized to the volume of the ROI.

### Spatial transcriptomics

Spatial transcriptomics on mouse brain tissue was performed as previously described ^71^. Briefly, fresh-frozen brains from WT and *Per1^-/-^* mice collected at ZT2 were embedded in OCT were cryo-sectioned at 10 µm thickness. Tissue optimization was performed according to the Visium Spatial Tissue Optimization User Guide (CG000238 Rev A, 10x Genomics). Tissue permeabilization of 18 minutes was identified as the optimal time following testing a range of incubation periods. For spatial library construction, we used Visium Spatial Gene Expression Slides (CG2000233, 10x Genomics) according to the Visium Spatial Gene Expression User Guide (CG000239 Rev D, 10x Genomics). Briefly, tissue sections were dried for 1 min at 37°C after sectioning, and then fixed in chilled 100% methanol for 30 min, followed by H&E staining (with hematoxylin for 5 min, followed by 2 min in eosin). Brightfield histology images were captured using a 20x objective on an Axio Z1 slide scanner (Zeiss). The library PCR was performed with 10 cycles and double-sided size selection with SPRIselect performed with a 0.55x and 0.7x ratio. Libraries were sequenced on the NextSeq550 (Illumina) using a High Output 150 cycle kit (Illumina) at the University of Queensland Sequencing Facility, with the read structure as: Read1 - 28bp, Index1 - 10bp, Index2 - 10bp, Read2 - 120bp. Each sample received more than 120 million reads.

### Spatial transcriptomics data analysis

The raw sequencing data in BCL format was converted to fastq format using bcl2fastq/2.17. The fastq reads were further trimmed to remove the remaining sequences of the template-switching oligos and the poly-A tails. The processed fastq reads were then mapped to the mouse reference genome (mm10 V3.0.0) using SpaceRanger V1.2.2 (10x Genomics). On average, in each sample, a spatial spot had more than 4000 genes detected. As a quality control step, spots with fewer than 200 genes and genes expressed in fewer than three spots were removed from the downstream analysis.

Genes differentially expressed in the SCN were identified through an integrative analysis that combined histological information with gene expression. The SCN region in each brain section was identified using high-resolution H&E staining and genetic markers commonly expressed in the SCN. Morphologically informed SCN annotations were used to select spatial spots within the SCN regions. For the spatially defined spots in the SCN, pseudo-bulk analysis was performed as outlined next. In each sample, the spots were randomly grouped into three pools of an equal sample size. The representative expression value for a gene in each pool was the average expression of that gene across the spots within the pool. Each pool was then considered as one replicate for the sample in the differential expression analysis, using a generalized linear model implemented in the edgeR package (Chen et al., 2020). The covariates were specified according to the animal for the design matrix of the linear model. Changes in gene expression were considered significant if the adjusted *p*-value was equal or less than 0.05.

### *In situ* hybridization

To further validate the differential spatial expression profiles of *Avp* and *Pcsk1n* detected by spatial transcriptomics, RNAscope *in situ* hybridization was performed on SCN slices of WT and *Per1^-/-^* mice using the RNAScope Multiplex Fluorescent Reagent Kit V2 (#323100, Advanced Cell Diagnostics). 10μm freshly frozen SCN slices were fixed in 4% paraformaldehyde at room temperature for an hour, followed by dehydration in gradient concentrations of ethanol. Brain sections were then incubated with protease reagent (Protease IV) at room temperature for 30 minutes. After washing in 1XPBS, the brain sections were treated with RNAscope probes, including *Vip*, *Avp*, and *Pcsk1n* (RNAscope Probe-Mm *Vip*; Cat No. 415961-T8, Mm *Avp*; Cat No 472261-T7, Mm *Pcsk1n*; Cat No 574741-T6; Advanced Cell Diagnostics) in the HybEZ Oven at 40°C for 2 hours to hybridize probes. The sections were then subjected to a three-step amplification with each amplification reagent at 40°C for 30 minutes each step, followed by incubation in RNAscope HiPlex Fluoro reagent at 40°C for 15 minutes to enable visualization of signals. Slides were counterstained with DAPI and cover-slipped using ProLong Gold Antifade Mountant (Cat no. P36930; Fisher Scientifics). Fluorescent RNA signals were captured using a super-resolution confocal microscope (Zeiss LSM900 Fast Airyscan 2, Carl Zeiss, Germany).

### Targeted Metabolomics

#### Sample preparation

Targeted metabolomics on liver samples from WT and *Per1^-/-^*mice before and after the 12-hour phase delay was used to investigate the effects of phase shifts on the liver metabolome in the absence of *Per1*. Briefly, fresh-frozen liver tissues were homogenized in MilliQ water (1:10; w/v ratio). Liver homogenates were used for the analyses of metabolites involved in energy metabolism (e.g., TCA cycle metabolites and Ketone bodies), fat metabolism (e.g., bile acids), microbiota metabolites (short chain fatty acids (SCFA)) and reactive carbonyls (especially biogenic aldehydes). Sample preparation for all panels included the addition of an internal standard solution (10µg/ml of 2-ethyl butyric acid (2EtBA) for short chain fatty acids, biogenic aldehydes, ketone bodies, and the TCA cycle panel and 1µg/ml of 1-(4-fluorobenzyl)-5-oxoproline (FOB) for bile acids panel) to sample homogenates. The procedures hereafter define further sample handling/processing and analysis protocol for the different metabolite panels.

#### Sample processing, instrumentation and data acquisition/analyses (bile acid panel 01)

Liver homogenates were deproteinized by adding ice-cold acetonitrile (100µL) containing 0.4% formic acid to 100µL homogenates. The resulting homogenates were vortexed, chilled at 4°C for 15 minutes, sonicated at high power at 15 °C for 10 minutes, and centrifuged at 10,000g at 4°C for 10 minutes for supernatant extraction. The supernatant was transferred to a 96-well extraction plate (Phree^TM^ Phospholipid) prewashed with 100% methanol, conditioned with 75% aqueous acetonitrile/0.2% formic acid. Using 2 to 3 psi vacuum pressure, the solution was slowly flushed through the resin of the extraction plate. The flow-through solution for each sample was collected into 2mL wells of the deep 96-well collection plate with additional washing using 0.4mL of the 75% acetonitrile/0.2% formic acid. Eluted solutions were transferred to 1.5mL microtubes and dried using a speed-vacuum concentrator at 35°C. Reconstitute the dried residue with an additional 50µL of the blank solvent (methanol: acetonitrile: water; 1:1:3), vortexed and transferred to HPLC vials with inserts for analysis. For quantification and analytical purposes, due to the unavailability of pure standards, a semiquantitative approach was utilized to calculate fold change across all samples. The peak area ratio of each analyte versus the FOB of the internal standard (IS), the concentration of IS, the volume of IS, and the weight of the tissue were used to normalize the values to the baseline and fold change calculations.

#### Sample processing, Instrumentation and Data acquisition/analysis for SCFAs, reactive carbonyls, the TCA Cycle and ketone bodies panel (panel 02)

The metabolites were quantified in their benzyloxyamide derivative form and 2-ethyl butyric acid (2EtBA) as an internal standard. Sample derivatization was achieved using O-benzylhydroxylamine (O-BHA) and N-(3-dimethylaminopropyl)-N-ethylcarbodiimide (EDC) ^72^. For samples, 100µL of tissue homogenates in milli-Q water was vortexed with 10µg/mL of 2EtBA in milli-Q water, followed by deproteinization by 200µL of ice-cold methanol solution. The resulting solution was vortexed and centrifuged at 14,000 rpm for 10 min at 4°C. The supernatant was incubated with 30µL of 0.1M O-BHA in methanol and 30µL of 0.25M of EDC in methanol at 25°C for 1 hour. After the incubation, the samples were diluted with 200µL of milli-Q water and extracted with 600µL of dichloromethane through vigorous shaking for 10 min followed by centrifugation 14,000 rpm for 5 min at 4°C. The tubes were tilted at an approximately 45-degree angle, and the upper aqueous layer was discarded along with any residual material. The organic DCM layer was evaporated to dryness using a nitrogen dryer set at 40°C. The residual was reconstituted in 50µL of 50% methanol, centrifuged at 14,000 rpm for 10 min at 20°C and the upper clear 50µL of supernatant was transferred to an HPLC vial containing inserts for LC-MS/MS analysis.

#### Instrumentation and data acquisition

For panel 01, the quantification of bile acids was achieved in an underivatized form ^73^. LC-MS/MS was performed using a QTRAP 5500 (AB SCIEX, Framingham, MA, USA) linear ion trap triple quadrupole LC-MS/MS mass spectrometer equipped with Turbo V electrospray ionization (ESI) source system united with Genius AB-3G nitrogen gas generator (Peak Scientific, Inchinnan, Scotland, UK) was used. The mass spectrometer was coupled with a Shimadzu LC30 series UPLC system (Shiamdzu Technologies) equipped with a degasser, a column over, a binary pump, and a temperature-controlled auto sampler. The temperature for the auto sampler was set at 10 °C, and the column over temperature was maintained at 45±1 °C. Chromatographic separation was implemented by the Water BEH C18 LC column (150 x 2.1mm, 130 Å, 1.7µm, Water, USA) under binary gradient conditions using mobile phase A (0.01% formic acid in LC grade milli-Q water) and mobile phase B (0.01% formic acid in acetonitrile) with a flow rate of 350 µL/min. Analytes were eluted using the binary gradient i.e., 25% mobile phase B from 0 - 0.01 min with a linear increase to 40 % from 0.01 to 12 min followed by a linear increase to 75 % till 14 min and 100 % until 16 min, hold at 100 % up till 22 min. Column washing and equilibration were achieved by a linear decrease to 2 % from 22 to 23 min, followed by 2% of mobile phase B for the next 3 minutes and equilibration using 25% B from 26.5 min to 30 min. The injection volume of 10 µL was set for method development and sample analysis. Relative levels of analytes were measured against the known concentration of the internal standard and the peak area ratio of the analyte to the internal standard. The mass spectrometer was operated in multiple reaction monitoring (MRM) scan mode to identify fragments in negative ionization mode at unit resolution. The detailed Q1 and Q3 transitions and optimized MS parameters have been summarized in Supplementary Table 1. The % CV of retention time and signal intensity of the internal standard across all samples was used as intra-run and inter-run quality control. Analyte software (version 1.7.1, AB SCIEX, Applied Biosystems Inc., USA) was used for system control and data acquisition. Experimental data processing and analysis were performed on MultiQuant software (version 2.0, AB SCIEX, USA).

For Panel 02, LC-MS/MS was performed similarly to what was described in previous studies ^72^. Briefly, an API 3200 (AB SCIEX, Framingham, MA, USA) triple quadrupole LC-MS/MS mass spectrometer equipped with Turbo V electrospray ionization (ESI) source system united with Genius AB-3G nitrogen gas generator (Peak Scientific, Inchinnan, Scotland, UK) was used. The mass spectrometer was coupled with an Agilent 1200 series HPLC system (Agilent Technologies) equipped with a degasser, a column over, a binary pump, and temperature-controlled auto sampler. The temperature of the auto sampler was set at 4 °C, and the column temperature was maintained at 25±1 °C. Chromatographic separation was implemented by Kinetex EVO C18 analytical column (100 x 2.1mm, 100 Å, 5µm, Phenomenex Inc., CA, USA) under binary gradient conditions using mobile phase A (0.1% formic acid in LC grade milli-Q water with 10mM ammonium formate; pH 3) and mobile phase B (0.1% formic acid in 9:1 of methanol: isopropanol solution) with a flow rate of 400 µL/min. Analytes were eluted using the binary gradient, specifically, 2 % mobile phase B from 0 - 0.5 min with a linear increase to 85% from 0.5 to 4 min followed by a linear increase to 99% to 8 min, held at 99 % to 13 min. Column washing and equilibration were achieved by a linear decrease to 1 % from 13.5 to 17 min, followed by 2 % of mobile phase B for the last 2 minutes. The injection volume of 5 µl was set for method development and sample analysis. Relative levels of derivatized analytes were measured against the known concentration of the internal standard and the peak area ratio of the analyte to the internal standard. The mass spectrometer was operated in multiple reaction monitoring (MRM) scan mode to identify fragments in positive ionization mode at unit resolution. The detailed Q1 and Q3 transitions and optimized MS parameters have been summarized in Supplementary Table 1. The % CV of retention time and signal intensity of the internal standard across all samples was used as intra-run and inter-run quality control. Analyte software (version 1.5.1, AB SCIEX, Applied Biosystems Inc., USA) was used for the system control and data acquisition. Experimental data processing and analysis were performed on MultiQuant software (version 2.0, AB SCIEX, USA).

### Glucocorticoid measurement

Mice were individually housed in cages equipped with wire grid floors for easy access and collection of samples without disturbing mice, and they had *ad libitum* access to food and water. Mice were acclimatized to the collection cages for 3 days prior to the first collection of feces to minimize the effects of stress on corticosterone levels. Mice were then subjected to the 12-hour phase delay, and fecal samples were collected at 4-hour intervals over a period of 24 hours under 12:12 LD before the 12-hour phase shift for baseline measurement and on the 3^rd^ day post shift to observe entrainment of corticosterone rhythm ^74^. The corticosterone metabolite was extracted and quantified using RIA (MP Biomedicals, USA) as previously described ^75^.

### Statistical analyses

Statistical analyses were performed with GraphPad Prism (GraphPad Software, California, USA). ClockLab analysis (Actimetrics, Illinois, USA) was used to analyze locomotor activity rhythms. Arrhythmicity of behavioral rhythms was determined using chi-square analysis on the ClockLab Analysis. GLMMcosinor analysis on R studio was used to statistically determine rhythmicity and acrophase of timeseries data (Parsons R, Jayasinghe O, White N, Rawashdeh O (2024), _GLMMcosinor: Fit a Cosinor Model Using a Generalised Mixed Modelling Framework, R package version 0.2.0.9000, https://docs.ropensci.org/GLMMcosinor/,https://github.com/ropensci/GLMMcosinor).

*CircaCompare* analysis method on R studio was applied to statistically compare differences in mesor, amplitude, and phase between rhythmic parameters including rhythms of vigilance states, locomotor behavior, and clock genes expression ^76^.

Metabolomics data was processed and analyzed using MetaboAnalyst (MetaboAnalyst 5.0, Xia Lab, Canada). All data were presented as mean ± SEM. Mann-Whitney U test, the student’s *t*-test, and two-way ANOVA with the Dunnett multiple comparison test was used; *p*-values equal or less than 0.05 were considered statistically significant.

## Acknowledgments

We thank Ms. Melanie Flint and Natalie Molotkov for technical assistance and Mr. Rex Parsons for statistical support. We are indebted to Dr. Steven Reppert and Dr. David Weaver for donating the mice and Dr. Horst W. Korf for sending the mice. We thank Dr. Brian Key for invaluable input on the manuscript and Kahli Jones for help with spatial transcriptomics.

## Author contributions

Conceptualization: OR and PK Methodology: PK, VK, TW, TV, IH and HO Resources: JB and TW

Investigation: PK, VK and TV Analysis: PK, VK, JB and SW Statistics: PK, NG and OJ Visualization: PK

Funding acquisition: OR and HO Supervision: OR Writing—original draft: PK

Writing—review & editing: PK, OR, JH, WI, IH, HO and GR

## Data and materials availability

All data are available in the main text and/or the supplementary materials.

