## Supplement Material for "A systemic clock brake: Period1 stabilizes the circadian network under environmental stress"

Pureum Kim *et al*.

**This file includes:**

Figs. S1 to S7

Fig. S1.

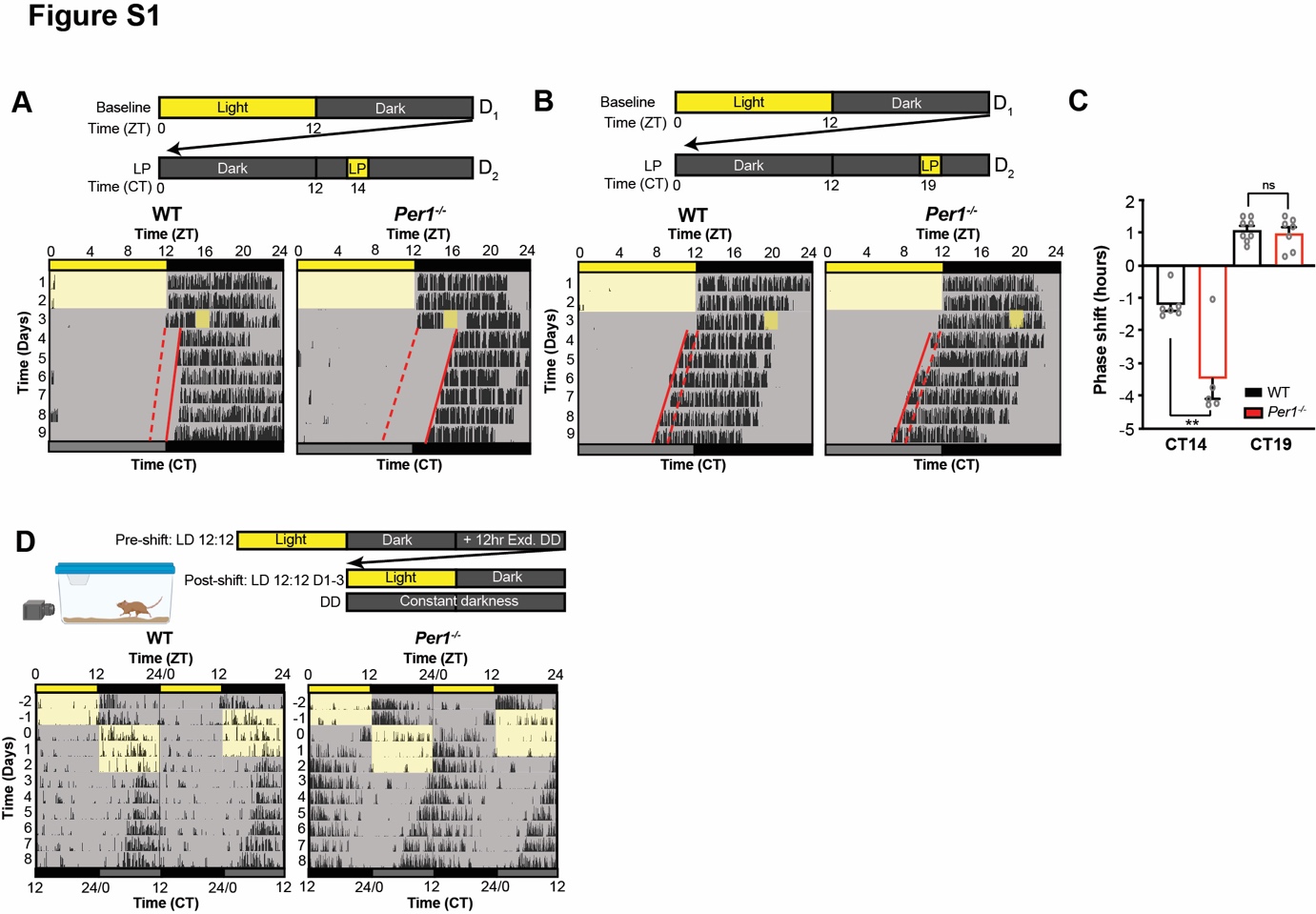

**Fig. S1.** (**A - B**) Representative actograms of locomotor activity of WT (Left) and *Per1^-/-^* (Right) mice in response to a light pulse at (**A**) CT14 or (**B**) CT19 (35 lux). (**C**) Quantification of the behavior phase shift to the light pulse in WT and *Per1^-/-^* mice (*p*=0.0008 for CT14 and *p*>0.05 for CT19, Student’s *t*-test, n=8 for WT and n=7 for *Per1^-/-^* mice). The data are presented as mean ± SEM. ** indicates *p*<0.005. (**D**) (Top) Experimental timeline: mice initially housed under LD 12:12 hours were phase-delayed by extending the dark phase by 12-hour and maintaining mice in the reversed LD for 3 days before subjecting the animals to constant darkness (DD). (Bottom) Double-plotted actograms of locomotor activity of WT and *Per1^-/-^* mice.

Fig. S2.

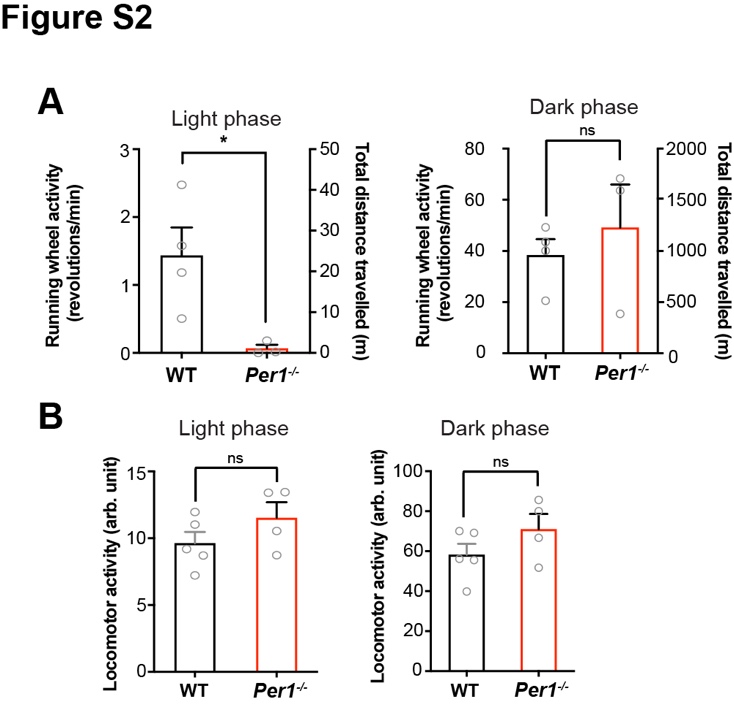

**Fig. S2.** (**A**) Quantification of running wheel activity during each phase in WT and *Per1^-/-^* mice (*p*=0.0379, Student’s *t*-test). (**B**) Quantification of locomotor activity recorded by infrared camera in WT and *Per1^-/-^* mice. Statistical significance is determined from comparison between different genotypes. The data are presented as mean ± SEM. * and ns indicate *p*<0.05 and *p*>0.05, respectively.

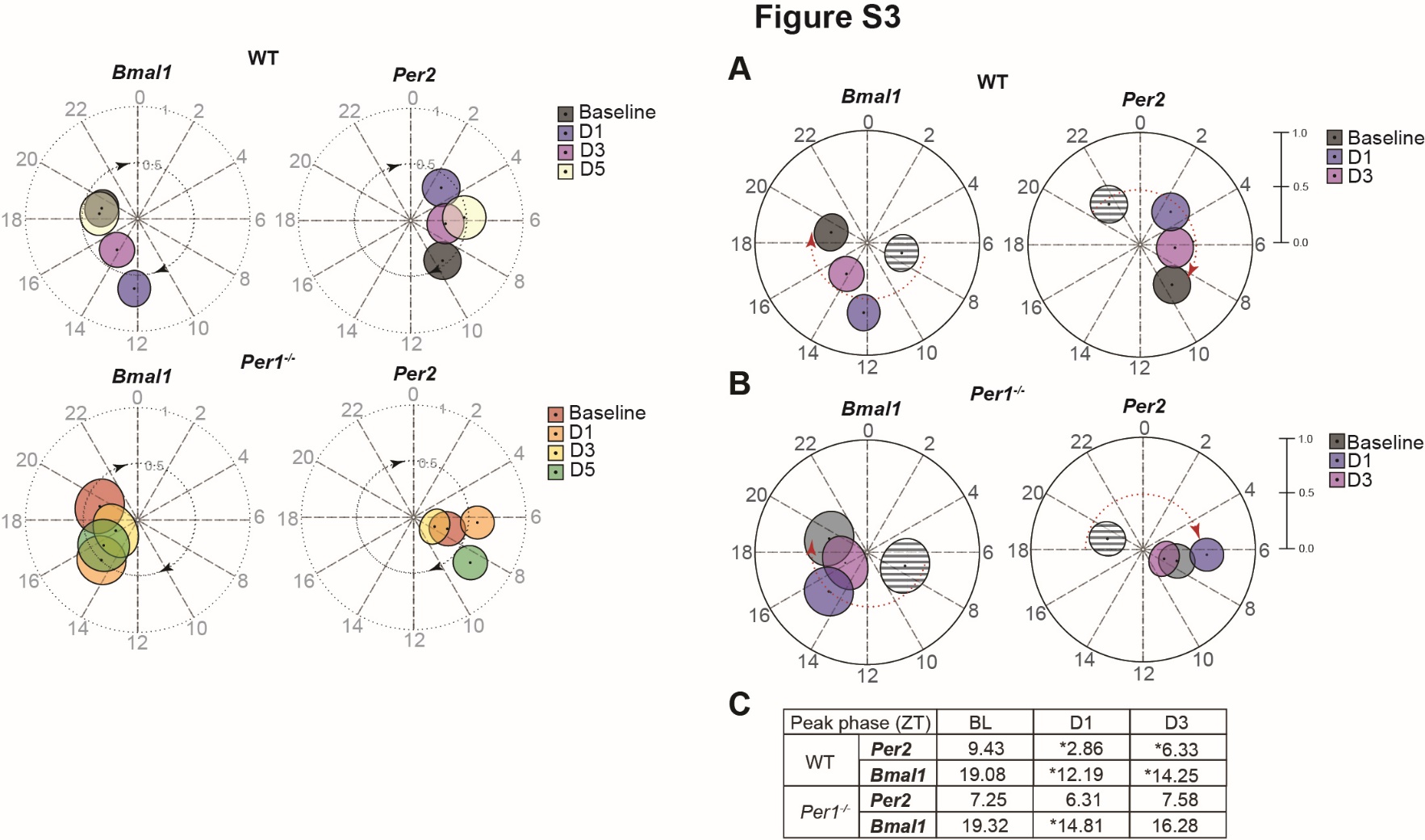
Fig. S3.

**Fig. S3. (A – B)** Re-entrainment of mRNA expression rhythms of *Bmal1* and *Per2* in the SCN on D1 and D3 post shift with corresponding baseline rhythm in in WT and *Per1^-/-^* mice. The peak phase in ZT time of expression rhythms is plotted as a circular shape. ZT time is indicated around the clock. The amplitude of expression rhythm is shown as the distance from the inner center to outer layers of circular grids. The arrow indicates the direction of re-entrainment following the phase shift. Faint grey circular shapes indicate the baseline based on the pre-shift light cycle. (**C**) Summary table showing the peak phases of the SCN *Bmal1* and *Per2* rhythms before and after the shift. Statistical significance (*p*<0.05) was determined by using *CircaCompare* and is indicated by an asterisk. N/A suggests that one or both gene expression profiles are arrhythmic.

**Fig. S4.**

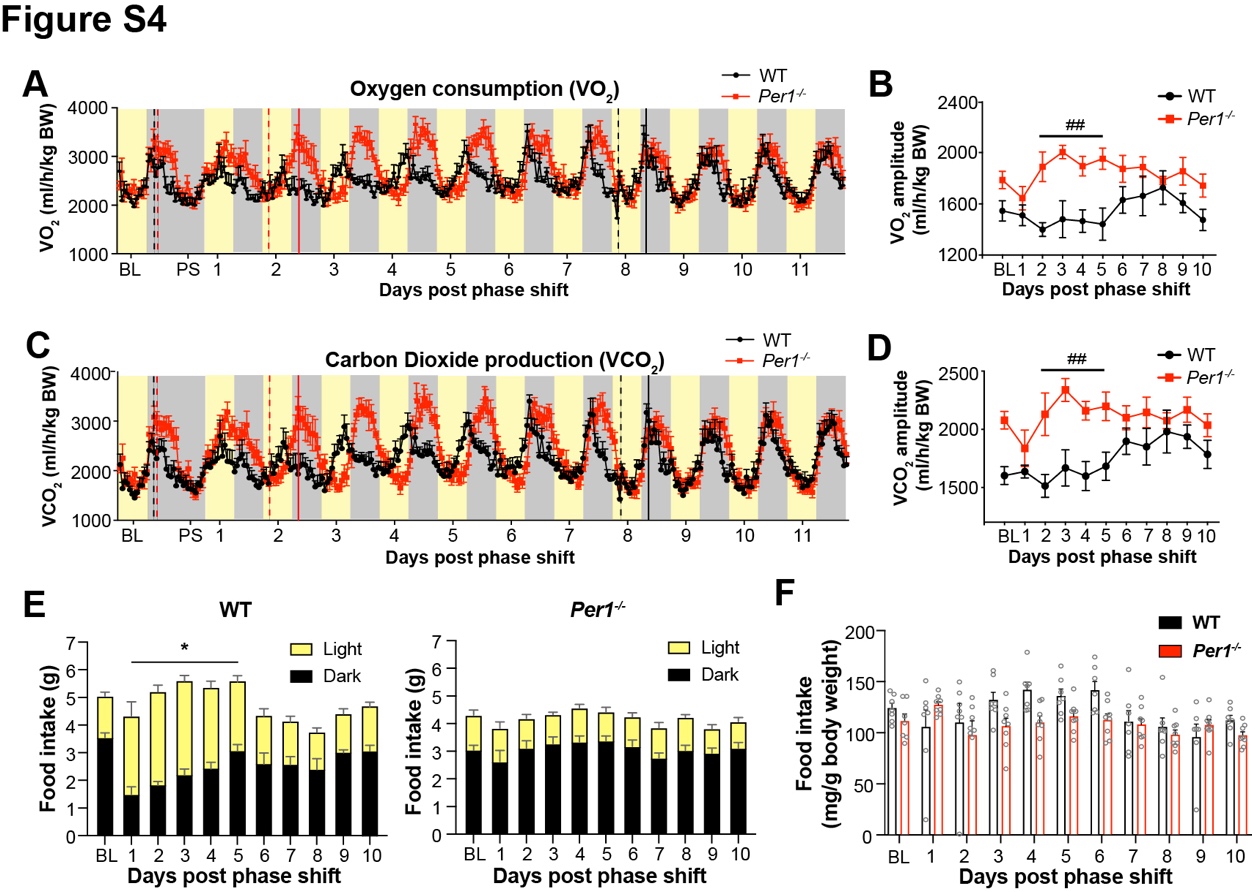

**Fig. S4. (A)** Rhythms of oxygen consumption (VO_2_) before and after the 12-hour phase delay in WT (black line) and *Per1^-/-^* (red line) mice. (**B**) Average amplitude of VO_2_ in WT and *Per1^-/-^* mice after the 12-hour phase delay (Effects of genotype *p*=0.0013, two-way ANOVA). (**C**) Rhythms of carbon dioxide production (VCO_2_) before and after the 12-hour phase delay in WT (black line) and *Per1^-/-^* (red line) mice. (**D**) Average amplitude of VCO_2_ in WT and *Per1^-/-^* mice after the 12-hour phase delay (Effects of genotype *p*=0.0015, two-way ANOVA). (**E**) Amount of food intake during each phase in WT (left) and *Per1^-/-^* (right) mice before and after the 12-hour phase delay (Effects of phase shift on food intake for both phases in WT *p*=0.0152, Two-way ANOVA). (**F**) Total daily food intake normalized to body weight for each genotype before and after the 12-hour phase delay. * indicates statistical significance for difference within each genotype and # indicates statistical significance for difference between genotypes. * and ** (##) indicate *p*<0.05 and *p*<0.005, respectively.

**Fig. S5.**

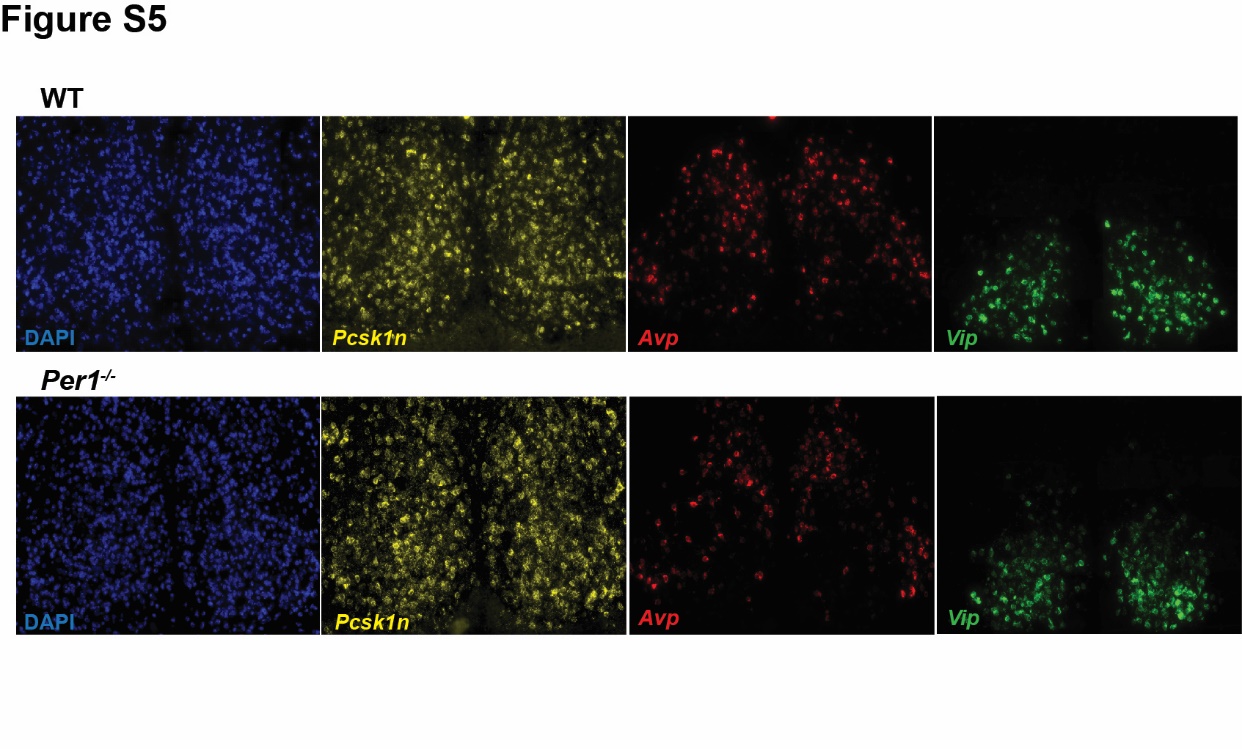

**Fig. S5.** Shown are the mRNA expression levels for the individual SCN neuropeptides (*Avp* and *Vip*) and the proprotein convertase subtilisin/kexin type 1 inhibitor (*Pcsk1n*). The brain slices shown here correspond to the WT and *Per1^-/-^* SCNs in Fig.5.

**Fig. S6.**

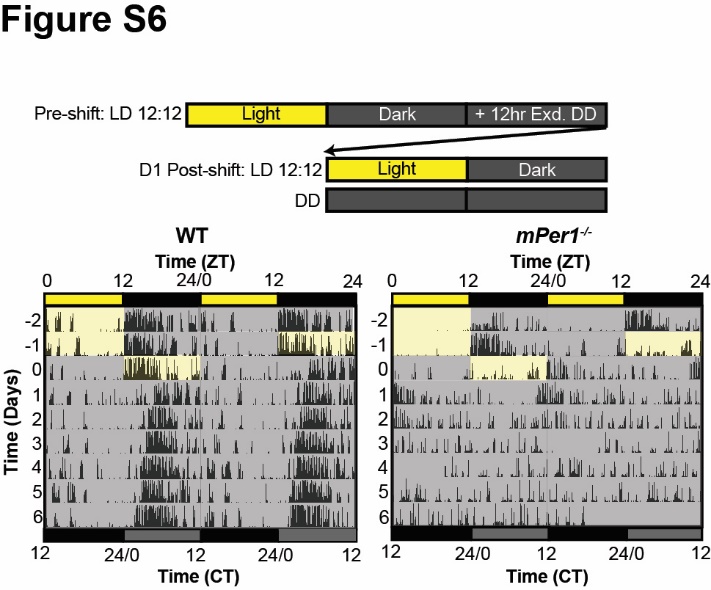

**Fig. S6.** (**A**) Experimental design (top) and representative actograms showing locomotor activity rhythms before and after the 12-hour phase delay in WT and *Per1^-/-^* mice (bottom).

**Fig. S7.**

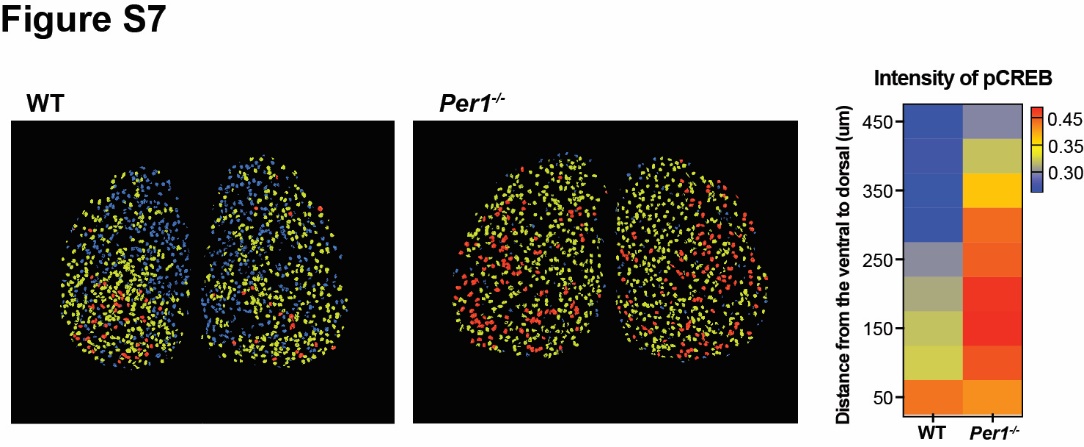

**Fig. S7. (Left)** The distribution of the pCREB immunofluorescence signal in the SCN of WT and *Per1^-/-^* mice. (**Right**) Heatmap showing the spatial distribution of different pCREB immunofluorescence signal intensity across the dorsoventral SCN axis.
